# The mechanism of motor inhibition by microtubule-associated proteins

**DOI:** 10.1101/2020.10.22.351346

**Authors:** Luke S Ferro, Lisa Eshun-Wilson, Mert Gölcük, Jonathan Fernandes, Teun Huijben, Eva Gerber, Amanda Jack, Katelyn Costa, Mert Gür, Qianglin Fang, Eva Nogales, Ahmet Yildiz

**Affiliations:** Department of Molecular and Cellular Biology, University of California, Berkeley CA; Department of Mechanical Engineering, Istanbul Technical University, Istanbul, Turkey; Department of Chemistry, University of California, Berkeley CA; Department of Imaging Science and Technology, Delft University of Technology, Delft, Netherlands; Biophysics Graduate Group, University of California, Berkeley CA; Press West Illustrations, Boston MA; Howard Hughes Medical Institute, Chevy Chase MD; Physics Department, University of California, Berkeley CA

**Keywords:** intracellular transport, microtubules, MAPs, kinesin, dynein

## Abstract

Microtubule (MT)-associated proteins (MAPs) regulate intracellular transport by selectively recruiting or excluding kinesin and dynein motors from MTs. We used single-molecule and cryo-electron imaging to determine the mechanism of MAP-motor interactions *in vitro*. Unexpectedly, we found that the regulatory role of a MAP cannot be predicted based on whether it overlaps with the motor binding site or forms liquid condensates on the MT. Although the MT binding domain (MTBD) of MAP7 overlaps with the kinesin-1 binding site, tethering of kinesin-1 by the MAP7 projection domain supersedes this inhibition and results in biphasic regulation of kinesin-1 motility. Conversely, the MTBD of tau inhibits dynein motility without overlapping with the dynein binding site or by forming tau islands on the MT. Our results indicate that MAPs sort intracellular cargos moving in both directions, as neither dynein nor kinesin can walk on a MAP-coated MT without favorably interacting with that MAP.

**HIGHLIGHTS:**

- MAP7 binds to a novel site and can coexist with tau on the MT.
- Kinesin-1 motility is biphasically regulated by MAP7 accumulation on the microtubule.
- MT decoration of MAPs inhibits motors even when they do not block the motor binding site.
- Motors need to interact with a MAP to walk on MAP-decorated MTs

## INTRODUCTION

Kinesin-1 (hereafter kinesin) and cytoplasmic dynein (hereafter dynein) are dimeric motors that transport organelles, vesicles, and nucleoprotein complexes to the plus- and minus-end of microtubules (MTs), respectively (Reck-Peterson et al., 2018; Verhey et al., 2011). Consistent with their fundamental roles in axonal transport and cell division, mutations that impair dynein- or kinesin-driven transport lead to multiple neurodegenerative and developmental disorders (Verhey et al., 2011). Live-cell imaging studies revealed that many cargos simultaneously recruit kinesin and dynein motors (Hancock, 2014). It remains unclear how cells regulate these opposing motors to achieve highly ordered and polarized transport of a wide variety of cargos. Recent studies indicated that signals that mediate polarized transport can be encoded on the MT tracks. MTs are formed by a combination of α and β tubulin isoforms and contain a large number of posttranslational modifications (PTMs). Distinct spatial patterns of this ‘tubulin code’ (Roll-Mecak, 2020) may sort the transport of intracellular cargos across the cell. Consistent with this model, mammalian dynein-dynactin exhibits a preference for binding tyrosinated MTs *in vitro* (McKenney et al., 2016). However, PTMs of tubulin only mildly affect the MT recruitment and velocity of kinesin motors (Kaul et al., 2014; Sirajuddin et al., 2014), suggesting that the tubulin code may not function as a binary switch of motor-driven transport.

The MT lattice is also decorated with a wide variety of structural MT-associated proteins (MAPs), which control the structural integrity and dynamics of MTs (Bodakuntla et al., 2019; Ramkumar et al., 2018). Cryo-electron microscopy (cryo-EM) has been used to visualize a number of MAPs on the MT surface, including tau and doublecortin (DCX), which bind across and stabilize longitudinal and inter-protofilament contacts, respectively (Kellogg et al., 2018; Manka and Moores, 2019). However, the MT footprint of many other MAPs remains to be determined. MAPs often extend from the MT surface through their disordered projection domain (Voter and Erickson, 1982), which can drive the formation of liquid condensates on the MT surface, as seen for tau (Siahaan et al., 2019; Tan et al., 2019). The transition of tau condensates to the solid state is the hallmark of Alzheimer’s and Parkinson’s disease (Wegmann et al., 2018).

MAPs exhibit distinct cellular localization in neurons, which appears to correlate with their regulatory role in the control of intracellular traffic (Bodakuntla et al., 2019; Gumy et al., 2017). For example, tau is concentrated in the axon and synapses of neurons (Wang and Mandelkow, 2016). Aberrant expression and phosphorylation of tau disrupt the transport of synaptic vesicles, while the knockdown of tau rescues defects in axonal transport in Alzheimer’s disease models (Ebneth et al., 1998; Ishihara et al., 1999). Despite differences in their domain organization and MT binding sites, MAPs are generally considered as obstacles for motor-driven transport. *In vitro* studies showed that kinesin-1 is inhibited by tau, DCX, MAP2, and MAP9 (Dixit et al., 2008; Monroy et al., 2018; Monroy et al., 2020), whereas dynein is inhibited by MAP2 and MAP9 (Lopez and Sheetz, 1993; Monroy et al., 2020). However, distinct MAPs can selectively recruit motors to MTs, as kinesin-1 and kinesin-3 driven transport depends on the presence of MAP7 and DCX, respectively (Barlan et al., 2013; Liu et al., 2012). *In vitro* studies showed that these MAPs activate motors through direct molecular interactions. MAP7’s projection domain interacts with the stalk of kinesin-1, whereas DCX and MAP9 interact with the kinesin-3 motor domain (Hooikaas et al., 2019; Liu et al., 2012; Monroy et al., 2018; Monroy et al., 2020).

Selective activation of kinesin motors by specific MAPs suggested that spatial patterning of distinct MAPs governs the recruitment and motility of motors in different cellular locations (Monroy et al., 2020). However, previous studies reported that dynein motility was not strongly inhibited by tau or MAP7 *in vitro* (Monroy et al., 2018; Tan et al., 2019), hence it remains unclear whether the “MAP code” model also applies to transport in the retrograde direction. Because dynein can efficiently avoid roadblocks on an MT due to its ability to take side steps (Ferro et al., 2019), it remained unclear whether MAPs can inhibit dynein as strongly as they inhibit kinesin.

The mechanism by which the MTBDs and projection domains of MAPs contribute to inhibition or activation of a motor is not well understood. Previous studies proposed that MTBDs of MAPs inhibit motors by overlapping with their binding sites (Monroy et al., 2020), and condensates formed by the projection domains could function as a selectivity barrier to control access of specific motors to the MT (Siahaan et al., 2019). It is also possible that MAPs specifically bind and recruit one motor type to the MT while inhibiting other motors. These models are not mutually exclusive, but their predictions have not been carefully tested *in vitro*.

In this study, we seek a mechanistic understanding of how MAP7, tau, and DCX affect kinesin and dynein motility *in vitro*. Using cryoEM, we determined the structure of MT-bound MAP7 and showed that the MT footprint of MAP7 overlaps with kinesin’s MT-binding site. While the MTBD of MAP7 inhibits kinesin motility, tethering of kinesin to the MAP7 projection domain supersedes this inhibition, resulting in a biphasic relationship between MT decoration by MAP7 and kinesin motility. The MT decoration by MAP7 also switches the direction of kinesin-dynein assemblies towards the MT plus-end due to selective activation and inhibition of these motors. Unlike MAP7, tau and DCX inhibit both motors regardless of binding site overlap. Molecular dynamics (MD) simulations provided a mechanistic explanation for how a MAP can inhibit a motor without a binding site overlap. Our results indicate that both anterograde and retrograde cargos would need to make favorable interactions with MAPs in order to walk along MAP-coated MTs.

## RESULTS

### MAP7 binds a novel site on the MT

We first sought to understand whether the inhibitory effect of a MAP on a motor could be explained by a binding site overlap model. Recent cryo-EM studies showed that tau binds on the outer-most crest of MT protofilaments and spans three tubulin monomers, whereas DCX binds between adjacent protofilaments and connects four tubulin dimers (Kellogg et al., 2018; Liu et al., 2012; Shigematsu et al., 2018). However, MAP7’s structure was unknown. Using cryo-EM, we visualized full-length MAP7 bound to the MT at 4.4 Å resolution (Figures 1A-C and S1A, Movie S1). MAP7 binds as an extended rod to a novel site on the MT surface and appears to straddle the inter-dimer and the intra-dimer tubulin interfaces on a protofilament. The binding site at the intra-dimer interface may be more flexible, since the density for MAP7 appears weaker over that tubulin interface (Figure 1C). The MAP7 projection domain and the E-hooks at the tubulin C-terminal tails were not visible in the cryo-EM map due to their conformational flexibility.

**Figure 1.**
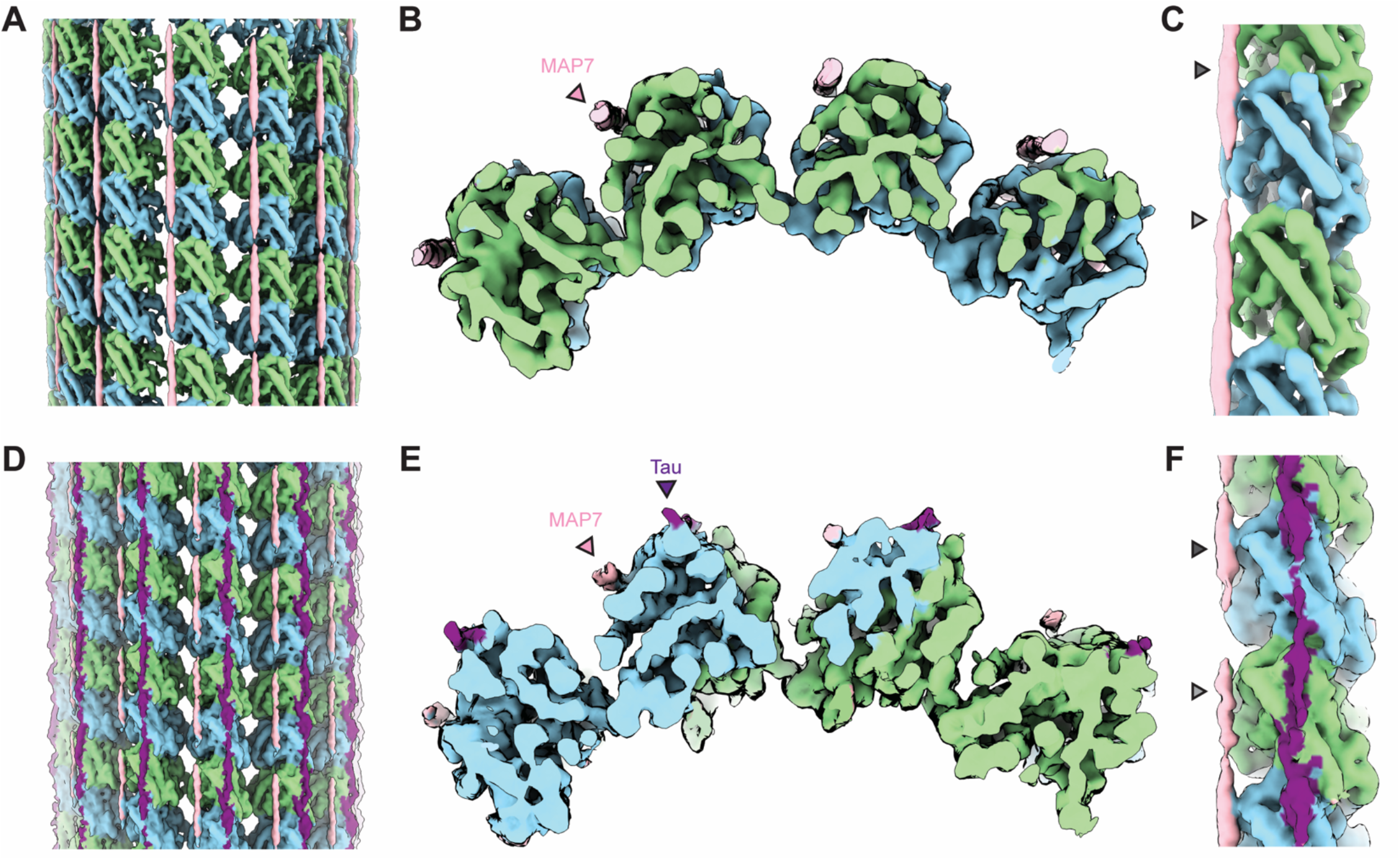
Cryo-EM reveals that MAP7 binds to a novel site on the MT. **(A)** Cryo-EM map of an MT decorated with MAP7; α-tubulin, β-tubulin, and MAP7 are shown in green, blue, and pink, respectively. MAP7 binds on the outer surface of the MT along protofilaments (PFs). The map is shown at a threshold of 3.2 in ChimeraX. **(B**) An end-on view of the MT wall showing the binding of MAP7. **(C)** Close-up of the structure highlighting the contact points of MAP7 with α - and β - tubulin (grey arrows). **(D)** Cryo-EM map of an MT co-decorated with MAP7-MTBD and tau, shown at a threshold of 2.1 in ChimeraX. **(E)** An end-on view of the MT wall showing the binding of the MAP7-MTBD and full-length tau. **(F)** Close-up of the MAP7-MTBD and tau co-decorated MT.

Because tau and MAP7 bind to different sites on the MT, we tested whether these MAPs can coexist on an MT. We obtained a 4.5 Å co-structure of MAP7’s MTBD (aa 60-170) with full-length tau on the MT (Fig. 1D-F and S1B). The costructure conclusively demonstrated that MAP7 and tau occupy different sites on the MT, suggesting that the removal of tau from MTs by MAP7 (Monroy et al., 2018) is not driven by a direct overlap between the MTBDs of these MAPs on the MT surface.

To determine whether the MAP7 binding site overlaps with motor binding sites, we superimposed previously determined structures of kinesin and dynein bound to the MT onto our MAP7 structure (Figure 2A). Surprisingly, there was a clear overlap between the MAP7 density with the binding sites of both kinesin and dynein at the intra-dimer interface between tubulin subunits (Lacey et al., 2019; Nishida et al., 2020; Redwine et al., 2012; Shang et al., 2014). In comparison, the MT-bound structure of tau (Kellogg et al., 2018) overlaps with the kinesin binding site, but not with the MTBD of dynein. Unlike these motors, the MT-bound structure of the N-terminal doublecortin domain of DCX (Manka and Moores, 2019) overlaps with neither kinesin nor dynein.

**Figure 2.**
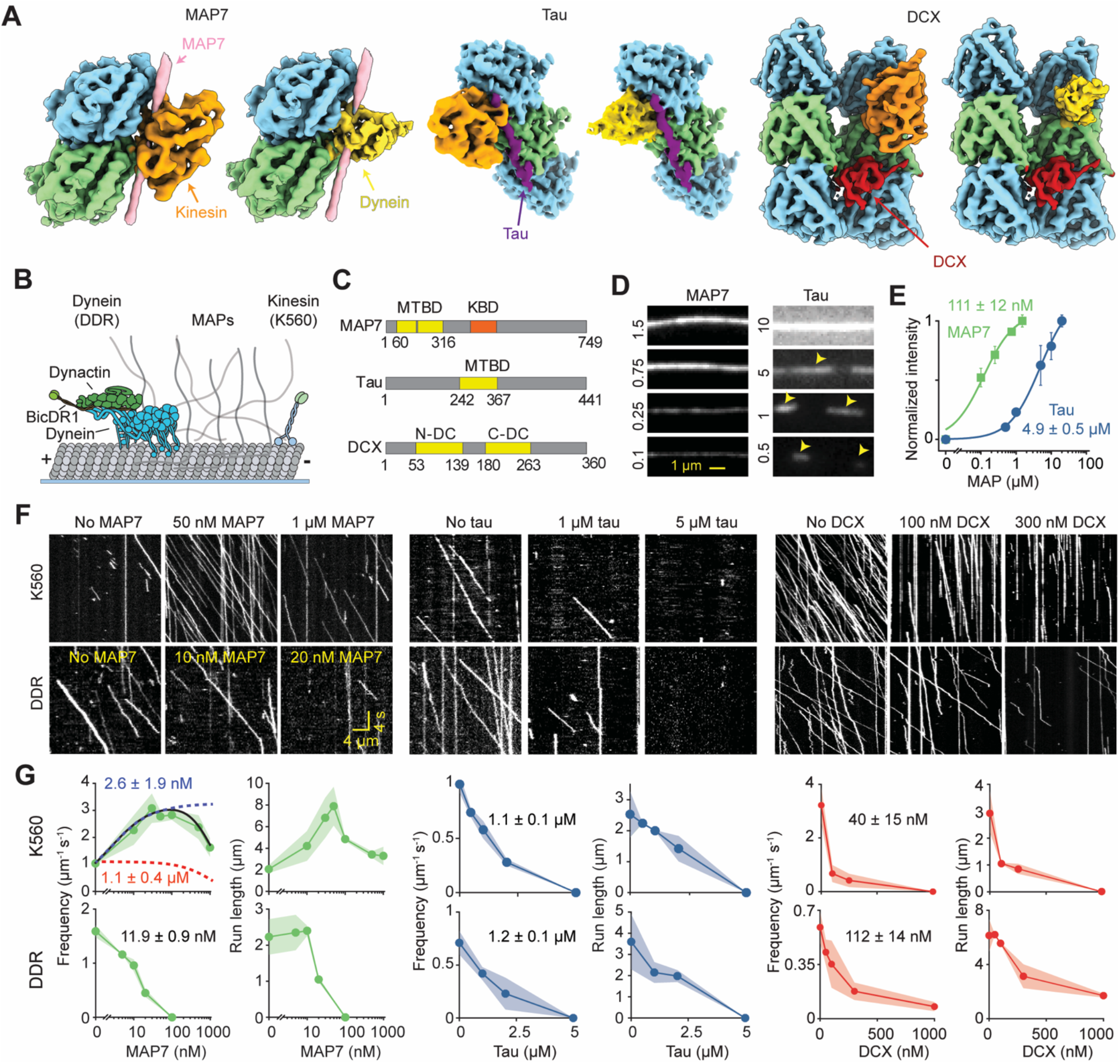
MAPs differentially regulate kinesin and dynein motility. **(A)** The density for MAP7, tau (EMDB ID: 7769), and DCX (EMDB ID: 1788) are superimposed with the kinesin motor domain (EMDB ID: 6353) and dynein MTBD (EMDB ID: 10060) bound to the MT. **(B)** Schematic of kinesin and dynein motility on MTs coated with MAPs. DDR assembled with mammalian dynein, dynactin, and BicDR1. Human kinesin-1 was truncated at its tail (K560). **(C)** Schematic of MAP7, tau, and DCX functional domains. DCX has an N-terminal and a C-terminal doublecortin domain (N-DC and C-DC, respectively) that bind to the MT. **(D)** Images of fluorescently labeled MAP7 and tau on single MTs. Higher concentrations (μM) were used for tau to account for its lower MT affinity. Yellow arrows point to tau islands at low concentrations. **(E)** Normalized intensity of MAP7 and tau binding to the MT (mean ± s.d.). K_D_ values (±s.e.) were calculated from a fit to the Hill equation (solid curves). From left to right, N = 54, 63, 58, 49, 73 for tau, and 44, 50, 68, 109 MTs for MAP7 (two technical replicates). **(F)** Representative kymographs of K560 and DDR motility in 150 mM KAc and 0.1% methylcellulose in the absence and presence of MAPs. **(G)** Run frequency and run length of K560 and DDR at different MAP concentrations (mean ± s.e.m.). The biphasic regulation of K560 run frequency with MAP7 was modeled by a fit to a Hill equation (solid black curve) to reveal the half-maximal activation (blue dashed curve) and inhibition (red dashed curve) concentrations (±s.e.). In other run frequency plots, IC_50_ values (± s.e.) were calculated from a fit with a single exponential decay function (not shown). For MAP7, N = 281, 463, 532, 836, 381, 433, 233 K560 motors, and 386, 235, 213, 146 DDR motors. For tau, N = 198, 160, 209, 99 K560 motors, and 185, 129, 100 DDR motors for tau. For DCX, N = 655, 187, 130, 0 K560 motors and 224, 237, 245, 104, 55 DDR motors (from left to right, two technical replicates).

### The regulatory role of a MAP cannot be predicted based on MT binding site overlap

The “steric overlap model” would predict that MAP7 inhibits both motors, tau inhibits only kinesin, and DCX inhibits neither motor (Figures 2A). To test these predictions, we decorated surface-immobilized MTs with full-length human MAP7, tau, or DCX and tested how these MAPs affect kinesin and dynein motility (Figure 2B-C and S2A-B). Consistent with previous reports (Siahaan et al., 2019; Tan et al., 2019), tau formed liquid condensates on surface-immobilized MTs at low concentrations (0.5 μM) and these condensates grew and merged at higher concentrations of tau (5 μM) under physiological salt (150 mM KAc) (Figures 2D and S2C). Unlike tau, MAP7 did not form liquid condensates and uniformly decorated the MT surface (Figure 2D) (Hooikaas et al., 2019). MAP7 bound the MT more strongly than tau, with a dissociation constant (K_D_) of 111 ± 12 nM, compared to 4.9 ± 0.5 μM for tau (Figure 2E). Although the MTBD of MAP7 can coexist with full-length tau on an MT (Figure 1E), full-length MAP7 inhibited tau binding to the MT surface with a half-maximal inhibition (IC_50_) of 270 ± 18 nM (Figure S2D-F), suggesting that MAP7 and tau compete on the MT through a mechanism other than steric binding site overlap (Monroy et al., 2018).

We next tested how these MAPs affect kinesin and dynein motility. Because full-length kinesin is autoinhibited by the folding of its tail onto the motor domain when not transporting a cargo (Friedman and Vale, 1999; Kelliher et al., 2018), we expressed constitutively active human KIF5B construct (K560, amino acids 1-560) that lacks the tail domain. Similar to kinesin, dynein is autoinhibited through specific interactions between its motor domains (Zhang et al., 2017), we assembled the dynein-dynactin-BicDR1 (DDR) complex that walks processively along MTs (Urnavicius et al., 2018) (Figure 2B). Previously, *in vitro* studies of MT motors have mostly been performed under low salt conditions, as motors either quickly detach from MTs or exhibit infrequent runs at physiological salt. Because the MT affinity of MAPs is also salt-dependent, MAP-motor interactions have been studied at nanomolar concentrations of MAPs, while cellular MAP concentrations are in the micromolar regime (Wegmann et al., 2018). We addressed this issue by adding 0.1% methylcellulose (a crowding agent), which enabled K560 and DDR motors to take extended processive runs in 150 mM KAc, with velocities of 829 ± 27 and 651 ± 100 nm s^-1^, respectively (mean ± s.d., Figure S3A). These conditions enabled us to investigate the mechanism by which motors are regulated by cellular MAP levels under physiological salt conditions.

MT decoration by tau reduced the run frequency and run length of K560 motors (Figure 2F, Movie S2), as previously reported (Dixit et al., 2008; Ebneth et al., 1998; Vershinin et al., 2007). Although DDR efficiently bypasses static obstacles on an MT due to its ability to take sidesteps (Ferro et al., 2019), tau inhibited DDR motility as strongly as it inhibited kinesin (Figure 2G). This finding is in contrast with a previous report that dynein is not inhibited by 0.5 nM tau in the absence of salt (Tan et al., 2019). We confirmed that tau was also inhibitory to both motors without added salt (Figure S3B-E, Movie S2), and both K560 and DDR quickly detached when they encountered a tau condensate on the MT (Siahaan et al., 2019) (Figure S3F). We also tested whether multiple motor assemblies walk more successfully on tau-coated MTs, as this was shown to be an effective strategy to bypass static obstacles on MTs (Ferro et al., 2019; Tjioe et al., 2019). MT decoration by tau fully inhibited the motility of beads driven by multiple K560 or DDR motors as strongly as it inhibited individual motors (Figure S4). Therefore, sidestepping and teaming up of motors are not effective strategies to bypass the high density of tau that naturally coats cellular MTs.

Unlike tau, decoration of MTs by 50 nM MAP7 increased K560 run frequency and run-length by 4-fold, as previously reported (Hooikaas et al., 2019; Monroy et al., 2020) (Figure 2F-G, Movie S2). However, K560 run frequency unexpectedly decreased as MAP7 reached its saturation concentration on the MT. The transition between the stimulatory and inhibitory effects occurred at near K_D_ of MAP7. In contrast to a previous report (Monroy et al., 2018), we found that run frequency and run-length of dynein sharply declined by the addition of MAP7 (Figure 2F). IC_50_ of DDR run frequency was 11.9 ± 0.9 nM MAP7 (±s.e.), a concentration 10-fold below the K_D_ of MAP7 (Figure 2G). For both K560 and DDR, velocity declined gradually as MAP7 concentration increased (Figure S5), likely due to crowding on the MT surface.

Although the main MT binding domain of DCX (Manka and Moores, 2019) does not overlap with kinesin or dynein (Figure 2A), motility assays showed that DCX strongly inhibits both K560 and DDR motility (Figures 2F-G and S5). Collectively, these results showed that tau and DCX inhibit both kinesin and dynein, whereas MAP7 inhibits dynein and regulates kinesin motility in a biphasic manner. Because kinesin can walk on MAP7-coated MTs despite an overlap, whereas tau and DCX inhibit dynein without an apparent overlap with the motor binding site, our results are inconsistent with the predictions of the binding site overlap model.

### Biphasic regulation of kinesin motility by MAP7

To understand the mechanism by which MAPs differentially regulate motors, we first investigated the nonlinear relationship between the extent of MT decoration by MAP7 and kinesin motility (Figure 3). Biphasic regulation arises when an enzyme is subjected to simultaneous activation and inhibition that dominate at different concentrations (Valenstein and Roll-Mecak, 2016). The fit of K560 run frequency under different MAP7 concentrations to a two-component Hill equation revealed that the half-maximal activation concentration was nearly 1,000-fold lower than the inhibition concentration (Figure 2G), enabling robust kinesin motility under a wide range of MAP7 concentrations. To understand the source of these opposing inputs, we first tested whether the biphasic effect was also observed with the full-length (FL) kinesin heavy chain (Figure 3A-B). Unlike K560, FL kinesin is autoinhibited and exhibits poor motility *in vitro* (Friedman and Vale, 1999) and MAP7 activates FL kinesin by interacting with its coiled-coil stalk (Figure 3A) (Barlan et al., 2013; Hooikaas et al., 2019; Tymanskyj et al., 2018). Consistent with these reports, we observed a substantial increase in the run frequency of FL kinesin in 50 nM MAP7 (Figure 3D). Similar to K560, the gains in run frequency and run-length decreased at higher MAP7 concentrations (Figure 3E), revealing a biphasic relationship between MAP7 and FL kinesin.

**Figure 3.**
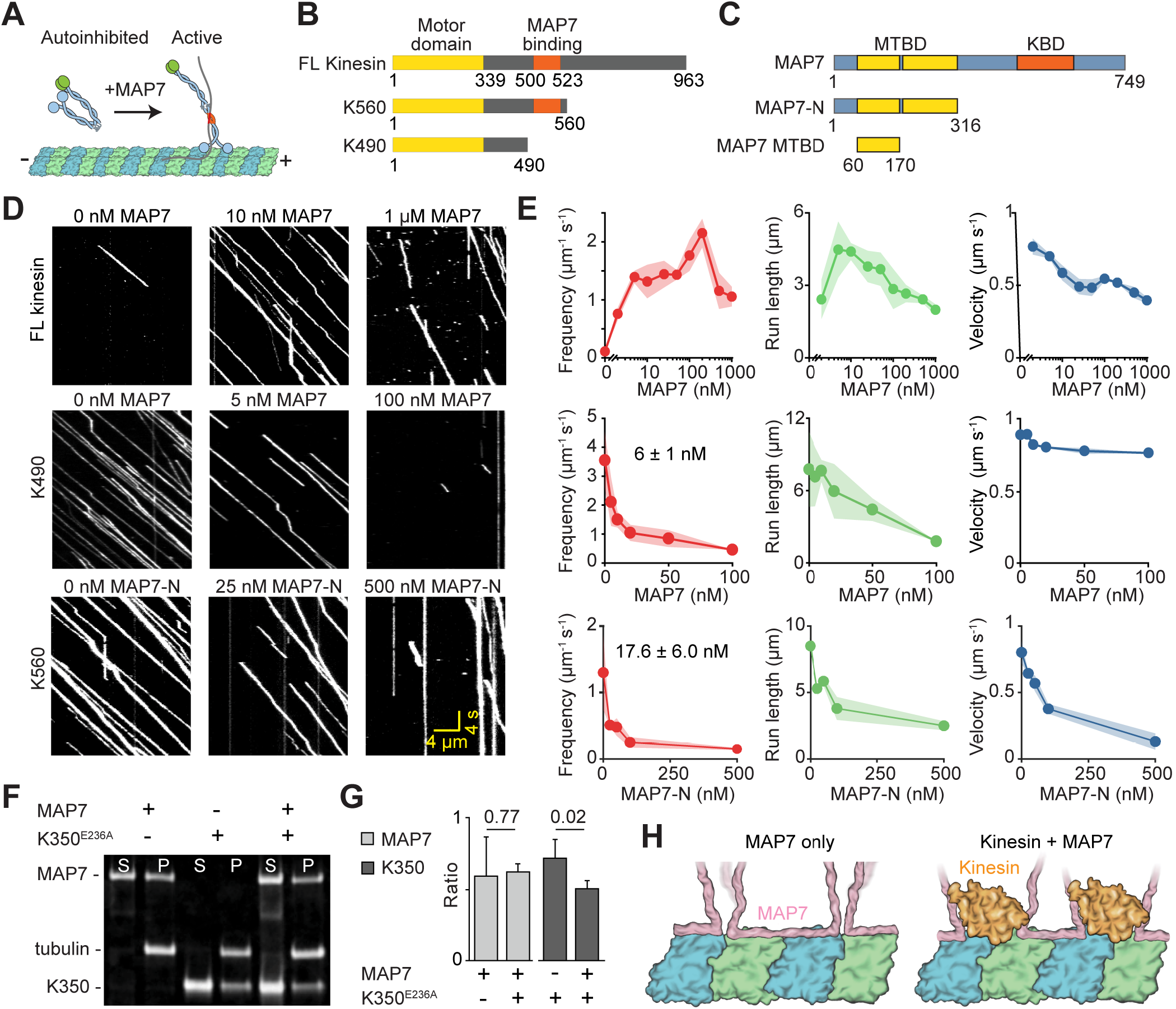
Biphasic regulation of kinesin by MAP7. **(A)** Schematic depicting the activation of kinesin by MAP7 on the MT. **(B)** Schematic of FL and tail-truncated kinesin constructs. K490 does not contain the MAP7 binding domain. **(C)** Schematic of full-length and truncated MAP7 constructs. **(D)** Kymographs show the motility of kinesin constructs in the absence and presence of full-length and truncated MAP7. Assays were performed in 150 mM KAc and 0.1% methylcellulose. **(E)** Run frequency, run length, and velocity of kinesin constructs under different MAP7 concentrations. Error bars represent s.d. For FL kinesin, N = 12, 658, 399, 527, 816, 664, 2434, 1303, 897, 802 runs from left to right. For K490, N = 855, 532, 581, 304, 457, 149 traces from left to right. For K560, N = 276, 168, 172, 121, 75 from left to right (two technical replicates). IC_50_ (±s.e.) of run frequency was calculated from a fit to a single exponential decay. **(F)** The denatured gel of MT pelleting assay tests MAP7 binding to MTs pre-decorated with K350^E236A^ in 150 mM KAc (S: supernatant, P: pellet). **(G)** Ratios of band intensities of (left) MAP7 to tubulin and (right) kinesin to tubulin in the pellet in G. Error bars represent s.d. between 3 technical replicates. p-values were derived from a two-tailed t-test. **(H)** A model for flexible rearrangement of MAP7 to accommodate kinesin on an MT.

Next, we tested a shorter kinesin construct (K490, amino acids 1-490) that lacks the MAP7-binding site (Figure 3B) (Monroy et al., 2018) (Monroy et al., 2018) (Monroy et al., 2018) (Monroy et al., 2018) (Monroy et al., 2018). We hypothesized that K490 would walk processively on MAP7-coated MTs, but its run frequency would not be enhanced. Surprisingly, we observed that full-length MAP7 inhibited K490 motility (IC_50_ = 6.0 ± 1.0 nM, ±s.e., Figure 3D-E, Movie S3), as strongly as it inhibited DDR (Figure 2). We also tested MAP7 constructs that lack the projection domain (MAP7-N: amino acids 1-316 and MAP7-MTBD: amino acids 60-170, Figures 3C and S6A) and thus are unable to bind kinesin. MAP7-N and MAP7 MTBD strongly inhibited both K560 and DDR motility (Figures 3D-E and S6B-C, Movie S4). We noticed that the degree of inhibition of K560 by MAP7-N (IC_50_ = 17.6 ± 6.0 nM) was comparable to inhibition of K490 by FL MAP7 (IC_50_ = 6 ± 1 nM, Figure 3E). Therefore, MAP7 fully inhibits kinesin motility in the absence of either the MAP7-interacting site on kinesin or the kinesin-binding site on MAP7. We concluded that the initial accumulation of MAP7 on the MT activates kinesin by tethering the motor to the MT via the projection domain, but the further accumulation of MAP7 reverses this effect due to the inhibition of kinesin motility by the MAP7 MTBD. Consistent with this conclusion, the addition of excess (1 μM) MAP7-N inhibits the motility of kinesin on MTs decorated with 50 nM FL-MAP7, whereas the addition of excess MAP7-C (amino acids 307-749), which lacks the MTBD but still interacts with K560, did not affect motility (Figure S6D-E).

The cryo-EM docking results also raise the question of how kinesin walks on MAP7-coated MTs when MAP7 is blocking its binding site at the intradimer interface. One possibility is that MAP7 possesses intrinsic flexibility at the intradimer interface (Figure 1C) to accommodate passing motors, as shown for DCX and kinesin-3 (Liu et al., 2012), and MAP4 and kinesin-1 (Shigematsu et al., 2018). To test this possibility, we performed MAP7 pelleting assays with MTs pre-coated with a rigor kinesin monomer (K350^E236A^, amino acids 1-350) in the presence of a nonhydrolyzable ATP analog, AMP-PNP. If MAP7’s binding site is truly blocked by the kinesin motor domain, MAP7 would not bind K350^E236A^-coated MTs. In contrast, we see that MAP7 binds the MTs in the presence of K350^E236A^ (Figure 3F-G), suggesting that MAP7 may rearrange on the MT to allow for kinesin to bind and walk along MTs (Figure 3H).

### MAP7 switches the directionality of cargos transported by kinesin and dynein

Given that MAP7 differentially regulates kinesin and dynein motility, we tested whether MAP7 could control the direction in which kinesin-dynein (KD) assemblies move along MTs (Chaudhary et al., 2019). We labeled FL kinesin and dynein with different fluorophores and connected FL kinesin to the BicDR1 subunit of the DDR complex via a GFP-nanobody linkage (Figure 4A). In the absence of MAP7, DDR walks processively on the MT whereas FL kinesin is autoinhibited (Figures 2 and 3). As a result, 80 ± 11% of KD assemblies moved towards the MT minus-end on undecorated MTs and their velocities were similar to the velocity of DDR complexes (Figure 4B-D). In the presence of 50 nM MAP7, DDR motility is inhibited while FL kinesin is activated (Figures 2 and 3). In this case, 93 ± 6% of KD assemblies moved towards the plus-end at velocities comparable to kinesin motors (Figure 4B-D, Movie S5). These assemblies move substantially faster than the outcome of a mechanical tug-of-war between active DDR and K560 motors assembled on a DNA scaffold (Belyy et al., 2016; Derr et al., 2012; Elshenawy et al., 2019). This is because the addition of MAP7 switches the activation state of opposite polarity motors and prevents motor tug-of-war that slows down transport. These results are in agreement with the view that MAPs can control the direction of cargo transport by activating one class of motor while inhibiting the other.

**Figure 4.**
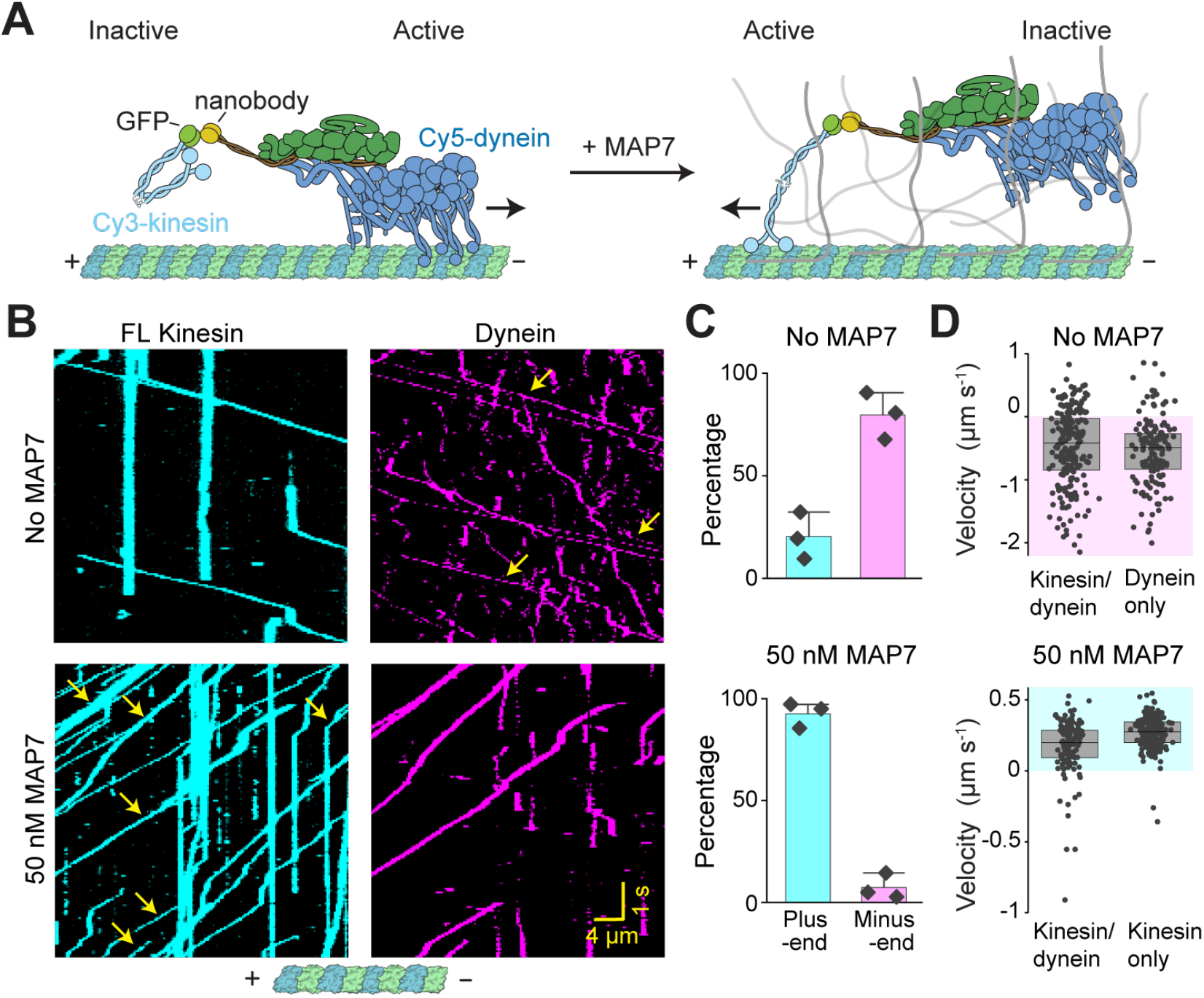
MAP7 switches the direction of kinesin-dynein assemblies on an MT. **(A)** Schematic of a model cargo assembled onto BicDR1. FL kinesin is attached to BicDR1 through a GFP-nanobody linkage. Dynein and FL kinesin are labeled with Cy5 and Cy3, respectively. In the absence of MAP7 (left), FL kinesin is inactive, whereas DDR is active. The addition of MAP7 switches the active motor (right). **(B)** Kymographs of kinesin-dynein assemblies in the absence and presence of 50 nM MAP7. Yellow arrows show the colocalization of Cy3-kinesin and Cy5-dynein trajectories. **(C)** The direction of complexes that contain both kinesin and dynein in the absence and presence of 50 nM MAP7. **(D)** (Top) The velocity of complexes that contain both kinesin and dynein versus dynein only in the absence of MAP7. N = 207, 151 from left to right. (Bottom) The velocity of complexes that contain both kinesin and dynein versus dynein only in 50 nM MAP7. N = 145, 435 from left to right. Experiments were repeated 3 times without additional KAc or methylcellulose.

### The MTBDs of MAPs inhibit motors

To understand how a MAP can inhibit a motor without overlapping with its MT binding site, we turned our attention to the regulation of motors by tau, which overlaps with kinesin, but not with dynein on the MT (Figure 2A). We first tested whether tau inhibits these motors through the formation of liquid condensates on the MT (Siahaan et al., 2019; Tan et al., 2019). We removed the projection domain by expressing a minimal tau construct containing its MTBD (Tau-MTBD, amino acids 242-367). Tau-MTBD only weakly interacted with MTs in 150 mM salt and bound to MTs at ~100-fold lower affinity than FL tau in the absence of added salt (Figures 5A-C). Unlike FL-tau, tau-MTBD did not form liquid condensates on MTs (Figure 5A), indicating that the projection domain is responsible for liquid-liquid phase separation (Tan et al., 2019) and helps to stabilize the MT binding of tau under physiological conditions. Importantly, tau-MTBD was sufficient to inhibit both K560 and DDR (Figure 5D-E), demonstrating that the projection domain and the formation of liquid condensates on an MT are not necessary for the inhibitory role of tau.

**Figure 5.**
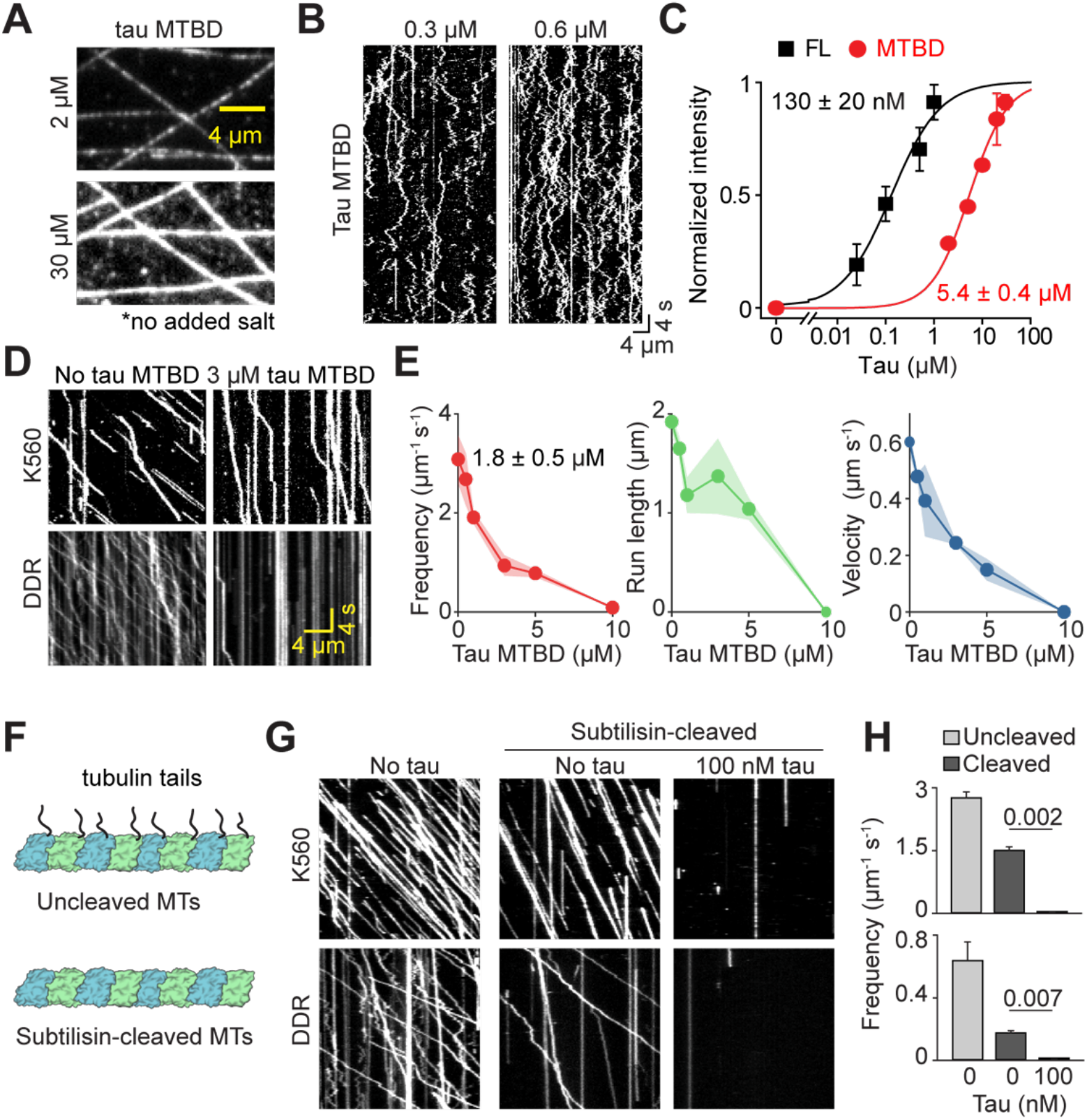
Tau inhibits kinesin and dynein in the absence of tau islands on the MT or the tubulin E-hooks. **(A)** Fluorescence images of tau MTBD decoration on MTs. **(B)** Diffusion of single tau MTBDs on the MT lattice. **(C)** The MT-binding affinity of FL tau and tau MTBD (N = 20 each condition, two technical replicates). Dissociation constants (±s.e.) were calculated from a fit to Hill Equation (black/red curves). **(D)** Kymographs of K560 and DDR motility in the absence and presence of tau MTBD. **(E)** The run frequency, run length, and velocity of K560 in different tau MTBD concentrations. N = 848, 801, 839, 568, 499, 46 from left to right. IC_50_ (±s.e.) of run frequency was calculated from a fit to a single exponential decay. **(F)** Schematic of subtilisin cleavage of the C-terminal tails of tubulin. **(G)** Kymographs of K560 and DDR motility on uncleaved and subtilisin-cleaved MTs in the presence and absence of 100 nM tau. **(H)** The run frequency (mean ± s.e.m.) of K560 and DDR motility on uncleaved and subtilisin-cleaved MTs in the presence and absence of 100 nM tau. p-values are calculated from a two-tailed t-test. Assays were performed in the absence of added salt and methylcellulose.

Previous studies showed that the negatively-charged tails of tubulin (E-hooks) capture motors through electrostatic attraction and retain them on the MT (Wang and Sheetz, 2000). To investigate whether tau inhibits motors by competing for these E-hooks, we cleaved the tubulin C-terminal tail by subtilisin treatment (Figure 5F). Consistent with previous reports (McKenney et al., 2014; McKenney et al., 2016; Sirajuddin et al., 2014), we observed a 50-70% reduction in the run frequency of kinesin and dynein on subtilisin-treated MTs (Figure 5G-H). Motors that landed on these MTs walked processively, but the addition of 100 nM tau fully abolished their motility in the absence of E-hooks (Figure 5G-H). We concluded that tau inhibits the binding of kinesin and dynein to the MT surface, rather than preventing their MT recruitment through the E-hooks.

### Tau sequesters tubulin residues required for dynein binding

Electrostatic interactions are critical for guiding motors to their tubulin binding sites (Li et al., 2016), where they form salt bridges with negatively-charged residues on the MT surface (Lacey et al., 2019; Redwine et al., 2012; Shang et al., 2014). The MTBD and regions that are proximal to MTBD of all three MAPs are highly enriched in polar and positively-charged residues (Figures 6A and S7A), which may disrupt these interactions and prevent the motor from stably binding to the MT. To test this possibility, we investigated the binding of dynein-MTBD to tubulin in the presence of tau by performing all-atom MD simulations. We used the cryo-EM structure of the R2 repeat of tau on tubulin as a template (Kellogg et al., 2018) to build an atomic model of dynein binding to a tubulin heterodimer in the presence of the R2 repeat (Figure S7B). Because flexible E-hooks could not be resolved by cryo-EM, they were not included in the atomic model. Structural studies revealed that docking of dynein to tubulin is driven by the rotation and translocation of a single helix (H1) towards the cleft between α and β tubulin, while the majority of the MTBD remains rigid upon binding to tubulin (Lacey et al., 2019). To monitor the docking of H1 to its stable binding site, we selected 25 conformers of dynein MTBD and a portion of the coiled-coil in the low-affinity state (Schmidt et al., 2015) and placed them ~1 nm away from the H1 docking site (Figure 6B). Starting from these unbound forms, dynein’s binding to tubulin was investigated by performing 250 ns long simulations for each system (12.5 μs in total). In the absence of tau, dynein formed stable salt bridges with the negatively-charged residues on the tubulin surface (Movie S6), as observed in MT-bound structures of dynein MTBD (Lacey et al., 2019). We observed that the formation of a salt bridge between K2996 of dynein-MTBD (K3298 in human dynein-1, Figure S7C) and E431 of β tubulin plays a major role in the docking of H1 to its stable binding position (Figure 6C-D). As a result, H1 was positioned within 0.5 nm root mean square distance (RMSD) from its docked position in 50% of the total simulation time and retained its expected angular orientation (Figures 6E and S7D-F).

**Figure 6.**
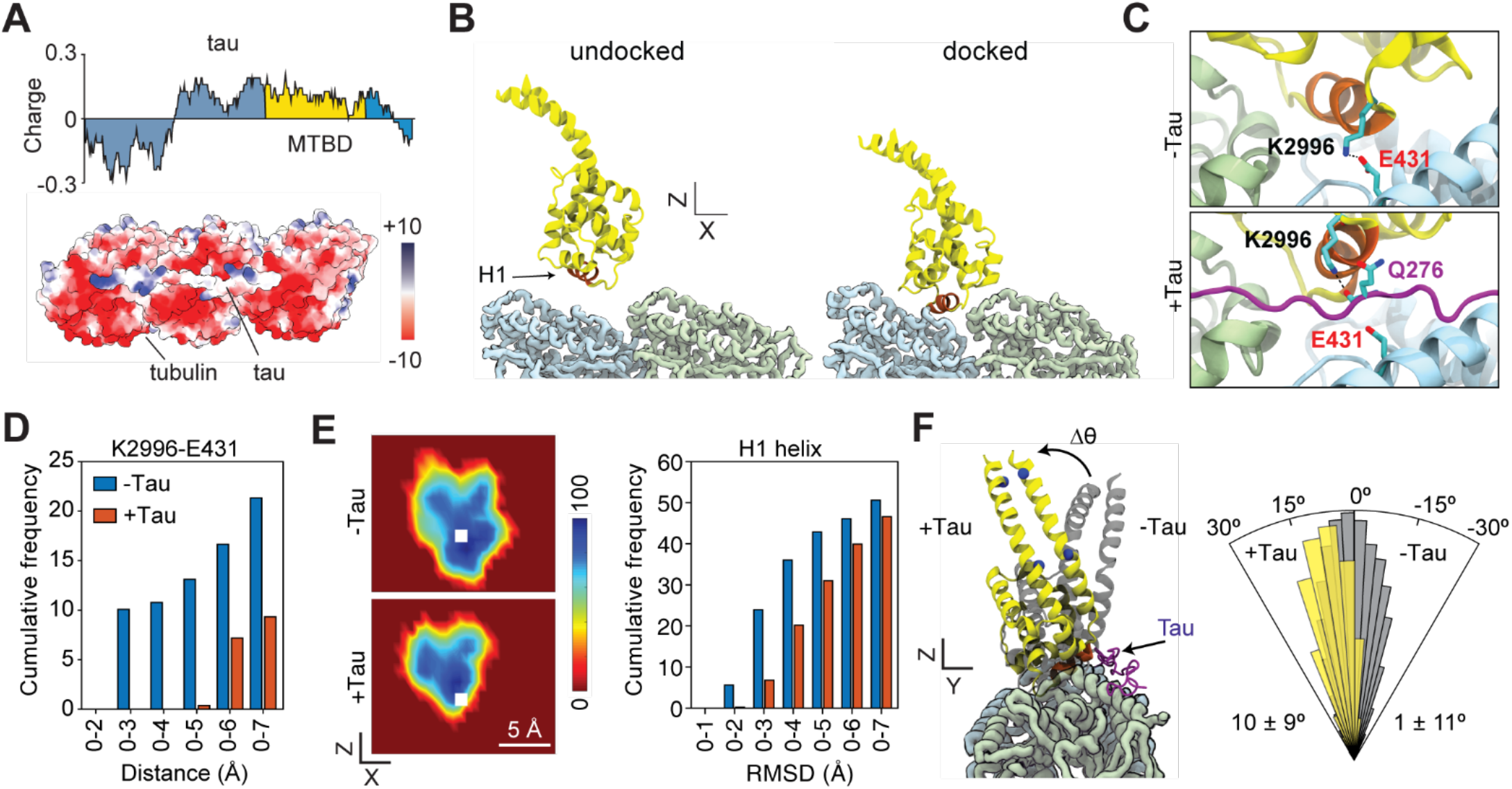
Inhibition of dynein motility by tau without an overlap on the MT. **(A)** (Top) Charge distribution on tau using a sliding window size of 30 amino acids. (Bottom) Electrostatic potential map of the MT surface and the R2 repeat sequence of tau (PDB: 6RZB). **(B)** MD simulations of dynein MTBD to a tubulin heterodimer. Dynein MTBD in the low-affinity state (PDB: 4RH7) was positioned 1.1 nm away from its tubulin binding site, and docking of dynein to the MT was visualized by all-atom MD simulations in the presence of solvent and ions. The H1 helix (red) rotates and moves towards the ridge between α and β tubulin, hence moving closer to the predicted H1 binding position in dynein’s high-affinity state (PDB: 6RZB) on the MT. **(C)** In the absence of tau, K2996 residue of dynein (K3298 in human dynein-1) forms a salt bridge with E431 of β tubulin to lower the H1 helix to its docking site. In the presence of tau, K2996 is positioned closer to tau and forms a hydrogen bond with the carbonyl oxygen Q276 (backbone) of tau (Q276). **(D)** Cumulative distribution of the distance between the basic nitrogen of K3298 of dynein and the acidic oxygens of E431 of β tubulin (left) with and without the R2 sequence of tau on the MT. **(E)** (Left) The heat maps show the sampling of H1 after it stably remains within 5 Å of its predicted position in the high-affinity state (white square) for 20 ns. (Right) The root-mean-square displacement (RMSD) of H1 from its predicted binding site. **(F)** (Left) Example snapshots of dynein MTBD in the presence and absence of tau were superimposed to show that dynein MTBD tilts away from tau and cannot retain its expected orientation on the MT. The vector that crosses the blue spheres of the crystal structure (PDB: 6RZB) was set to 0°. (Right) The angular orientation (Δθ) histogram of the stalk coiled coils in the presence and absence of tau (mean ± s.d.).

We next performed these simulations in the presence of the R2 peptide of tau on tubulin. We did not observe a steric clash between dynein’s MTBD and the tau backbone, and dynein was able to form salt bridges with negatively-charged residues of tubulin. However, the salt bridge between K2996 of dynein and E431 of β tubulin could not be formed due to hydrogen bonding between K2996 and the carbonyl oxygen of a well-conserved polar side chain (Q276) of the tau backbone (Figure 6C-D, Movie S6). As a result, the presence of tau on tubulin prevented H1 from moving near to its docking site (Figures 6E and S7D) and attaining correct angular orientation upon MT binding (Figure S7E-F). Dynein was also tilted ~10° away from tau compared to its high-affinity conformation on tubulin (Figure 6F, Movie S7). Collectively, our simulations indicated that tau does not prohibit transient interactions between dynein and the MT surface, but prevents dynein MTBD to switch to its high-affinity conformation even though it does not overlap with dynein on the MT.

## DISCUSSION

Kinesin and dynein motors deliver critical cargos to many destinations far from the cell center, but it still remains unclear how these motors walk on MTs that are densely decorated with structural MAPs. *In vitro* reconstitutions of kinesin and dynein motility on purified MTs have contributed significantly to our understanding of how these motors are regulated by specific MAPs. However, the mechanism by which MAPs inhibit or activate motors remains elusive. Do MAPs inhibit motors by overlapping with their MT binding sites? Do they selectively recruit motors by forming liquid condensates on the MT? How can a motor overcome inhibitory inputs by a MAP and transport its cargo? Using a combination of protein engineering, cryo-EM imaging, and MD simulations, we tested how steric overlap between the MTBDs of MAPs and motors on the MT, and how the projection domains of MAPs contribute to this regulation. Our results challenge previous views of motor regulation by MAPs and provide a broad framework for how MAPs specifically regulate motor proteins.

### Understanding the biphasic regulation of kinesin by MAP7

While MAP7 activates kinesin-1 on MTs at low concentrations (Hooikaas et al., 2019), we found that it inhibits motility at higher concentrations, leading to biphasic regulation of kinesin-1 motility. Based on our observations and previous reports, we propose a possible mechanism for this regulation. The MAP7 projection domain interacts with the coiled-coil stalk of kinesin-1 (Monroy et al., 2018), which rescues kinesin from autoinhibition and initiates processive motility along the MT (Hooikaas et al., 2019). Due to the weak affinity of kinesin-1 for MAP7 (K_D_>10 μM)(Hooikaas et al., 2019), the motor can disrupt its interaction with one MAP while forming a favorable interaction with another MAP as it walks through the dense “brush” of MAP projections on the MT surface. These interactions prevent the motor from diffusing away after it transiently dissociates from the MT. As a result, the motor quickly rebinds and continues to walk from the same location on the MT lattice, increasing its apparent run length. When the MT surface is nearly saturated with MAP7, run frequency and run length of kinesin are reduced because the inhibitory effect of MAP7’s MTBD begins to be more dominant than the activation from the projection domain. Kinesin and MAP7 have overlapping binding sites on the MT, and thus kinesin may have to wait for MAP7 to detach before taking a step. It is also possible that MAP7 rearranges on the MT surface to allow for motor binding, as seen with DCX and MAP4 (Liu et al., 2012; Shigematsu et al., 2018). Together, activation of motility by MAP7’s projection domain and inhibition by MAP7’s MTBD combine to yield a biphasic effect on kinesin motility. These results indicate that modulation of MAP7 accumulation on cellular MTs could tune kinesin-1 activity along these tracks.

### Inhibition of motors by MAPs

We find that the inhibitory role of MAP7 and tau on kinesin and dynein motility is primarily driven by their MTBDs. This result was unexpected because the two MAPs bind in different locations on the MT surface (Figure 1). MAP7-MTBD overlaps with the kinesin and dynein binding sites and may prevent these motors from binding the MT. However, the MTBD of tau inhibits dynein motility without a binding site overlap or liquid-liquid phase separation on the MT (Figure 7B). We also observed that both kinesin and dynein were inhibited by DCX, which has no overlap with these motors on the MT (Manka and Moores, 2019). Together, our data suggest that the effect of a MAP on a motor cannot be predicted based on the presence or absence of a binding site overlap.

**Figure 7.**
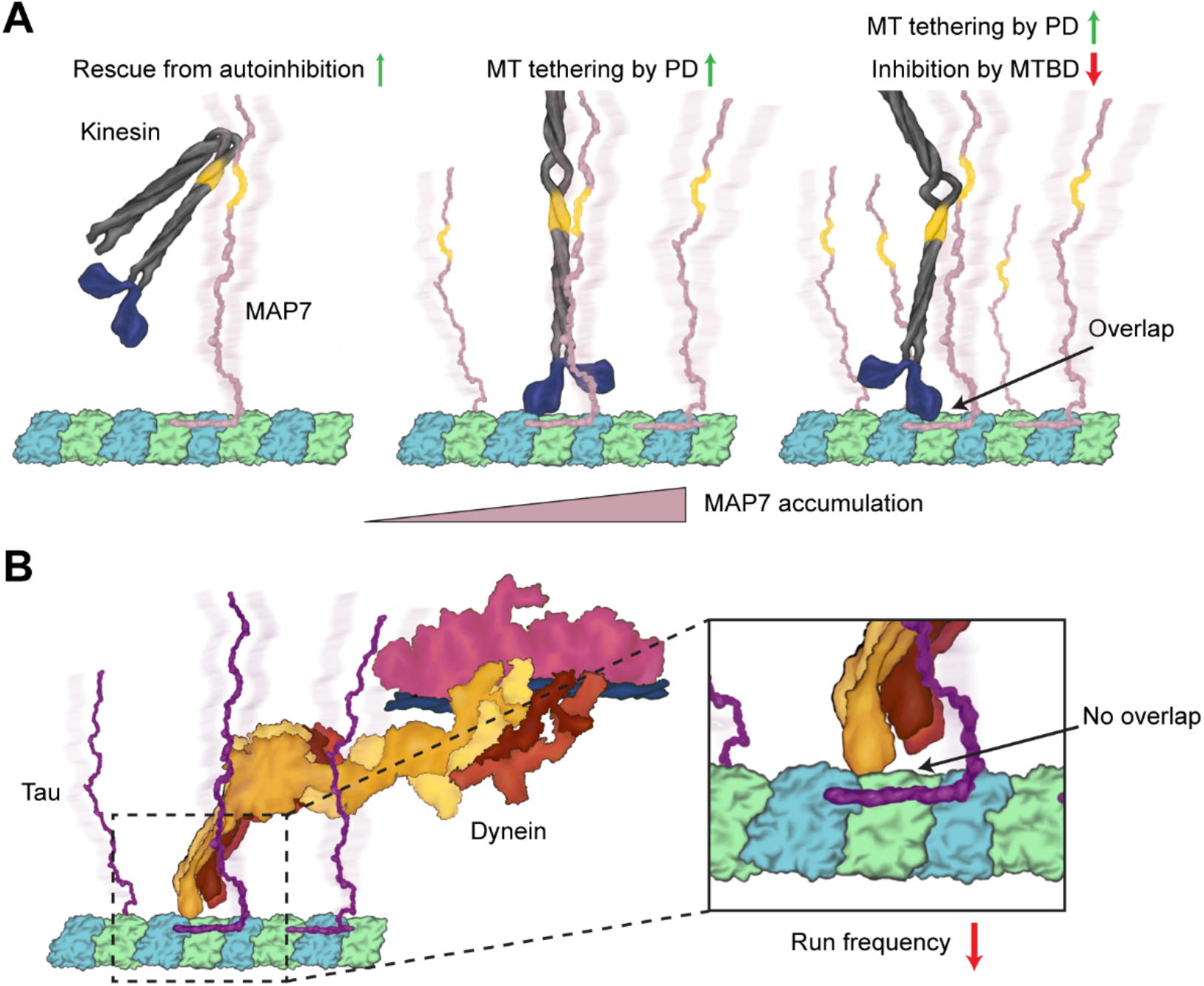
Model for regulation of MT motors by MAPs. **(A)** (Left) Kinesin is auto-inhibited in solution. Binding of kinesin to MAP7’s projection domain relieves the motor from auto-inhibited conformation. (Middle) Interaction between kinesin’s stalk and MAP7 projection domain (PD) serves as a tether that prevents the dissociation of kinesin from MT. Kinesin walks along MAP7-coated MT even though it competes for the same binding site with MAP7 MTBD. (Right) As the MT surface is nearly saturated with MAP7, MTBD inhibits kinesin motility either through steric overlap or electrostatic repulsion between the positive charges. **(B)** Unlike the kinesin-MAP7 pair, dynein does not specifically interact with tau’s projection domain. In this case, positive charges on tau and dynein MTBD repel each other and compete for the negatively charged residues on the MT. As a rest, tau inhibits dynein motility without overlapping between their MT binding sites.

Our MD simulations provide an explanation for how MAPs prevent motors from walking along MTs without a binding site overlap (Figure 6). The MT surface is strongly negatively charged, which favors the binding of the positively charged motor domains of motors (Li et al., 2016). The MTBDs of MAPs contain polar and positively charged residues and their MT binding can screen some of the critical salt bridges between the motor and the MT surface, as we observed in MD simulations of tau and dynein on tubulin. These interactions can inhibit the stable binding of motors to the MT and accelerate their dissociation as they step along the MT. In addition to the MAP/tubulin interface, negatively charged tubulin C-terminal tails may interact with the proximal regions of MAP MTBDs, which are strongly positively charged. While we observed that MTBDs of MAPs are still inhibitory to motors in the absence of tubulin tails, future work will be required to test whether PTMs of tubulin tails affect the interactions between MAPs and motors.

### Motors must interact with a MAP to walk on MAP-coated MTs

Previous studies showed that MAPs are largely inhibitory to kinesin motors *in vitro*; exceptions include MAP7, which activates kinesin-1, as well as DCX and MAP9, which activate kinesin-3 (Hooikaas et al., 2019; Liu et al., 2012; Monroy et al., 2018; Monroy et al., 2020). In these cases, the motor specifically interacts with the MAP, suggesting that MAPs inhibit kinesins unless the motor makes a favorable interaction with the MAP (Monroy et al., 2020). However, this idea had previously not been tested directly. Consistent with this view, we observed that tau and DCX inhibit kinesin-1 motility, while MAP7 specifically recruits and activates this motor on the MT (Hooikaas et al., 2019; Monroy et al., 2020). We showed that kinesin needs to interact with MAP7’s projection domain to overcome the negative effect of MAP7 binding to the MT. The MAP code model was not fully applicable to dynein, as dynein was reported to walk on MTs decorated with tau, MAP7, or DCX even though it does not form any known interactions with these MAPs (Monroy et al., 2018; Monroy et al., 2020; Tan et al., 2019). In contrast to these reports, we found that dynein is strongly inhibited by higher concentrations of these MAPs under physiological salt. These results strongly support the MAP code model, as neither dynein nor kinesin can walk on a MAP-coated MT without forming favorable interactions with that MAP.

### Differential regulation of motor activity by MAPs

In cells, the destination of specific cargos may be encoded by modifications of the MT tracks. In comparison to PTMs of tubulin (Kaul et al., 2014), MAPs have stronger effects on motors, suggesting that a “MAP code” on MT tracks may regulate bidirectional transport along MTs. This view has been supported by the observations that MAPs that localize to dendrites favor kinesin-3 motility on MTs *in vitro* and loss of these MAPs impede kinesin-3 driven transport in dendrites (Liu et al., 2012; Monroy et al., 2020). Similarly, the loss of MAP7 prevents kinesin-1 recruitment to MTs in human cells (Hooikaas et al., 2019). Consistent with the MAP code model, the addition of MAP7 causes a shift in the direction of isolated phagosomes on MTs *in vitro*, and this has been attributed to an increase in the MT-binding rate of kinesin-1 (Chaudhary et al., 2019). Similarly, we observed that the addition of MAP7 almost fully switches the direction of kinesin-dynein assemblies from the minus-end to the plus-end, favoring kinesin-driven transport. This major shift in direction is due to reciprocal activation and inactivation of kinesin and dynein by MAP7, rather than accelerated rebinding of kinesin-1 to MTs.

*In vitro* studies showed that interactions between kinesin-1 and MAP7 are not sufficient to overcome the negative interactions with other MAPs (Monroy et al., 2020). Therefore, it remains unclear how motors transport cargos in the presence of other inhibitory MAPs in cells. In neurons, the spatial patterns of MAP localizations across the cell may play a major role in sorting cargos transported by specific motors. Tau is concentrated in the axon (Monroy et al., 2020), DCX is enriched at the ends of the dendritic and axonal extensions while MAP7 is found throughout the neuron (Ramkumar et al., 2018). Similarly, tubulin PTMs are not uniformly distributed on adjacent MTs, and a subset of these MTs contains singular or a subset of these modifications (Tas et al., 2017). Super-resolution imaging studies in cells are required to test whether PTMs of tubulin regulate the recruitment of specific MAPs to MTs and whether there are MTs without MAPs or with a singular MAP to allow motors to transport cargos along these tracks.

While *in vitro* studies identified MAP7 and DCX as activators of kinesin-1 and kinesin-3 (Hooikaas et al., 2019; Liu et al., 2012; Monroy et al., 2018; Monroy et al., 2020), respectively, positive interactions between dynein and a specific MAP have not been reported, and its motility is inhibited by all three MAPs tested in this study. Although there are nearly 40 members of the kinesin motor family that bind specific cargos and localize to different regions in human cells, there is a single isoform of dynein responsible for nearly all minus-end directed transport in the cytoplasm. Therefore, the dynein transport machinery may possess a different mechanism to distinguish between axonal versus dendritic cargos. Dynein is recruited to specific cargos by coiled-coil adaptor proteins (Canty and Yildiz, 2020). Some of these adaptors, such as JIP1 (Fu and Holzbaur, 2013), contain an SH3 domain that is known to interact with proline-rich regions (Li et al., 2012). Tau, MAP7, DCX, and other MAPs have proline-rich regions projecting from the MTBD (Goode et al., 1997; Hooikaas et al., 2019; Liu et al., 2012; Manka and Moores, 2019), raising the possibility that regulation of the retrograde transport by MAPs is driven by cargo adaptor proteins, rather than dynein itself. Future studies are required to test whether this model can be generalized for the regulation of other cargo adaptors with specific MAPs.

## ACKNOWLEDGEMENTS

We thank Andrew Carter for helpful discussions, Laura Nocka for SEC-MALS experiments, Ben LaFrance and Joe Atherton for MT processing in RELION, Paul Sauer for MT pelleting assays, Daniel Toso and Abhiram Chintangal for microscopy and computational support, Julia Torvi for the pelleting assays, the QB3 Macro lab for competent cell lines and TEV protease purification, the UC Berkeley Cell Culture Facility for providing the insect cells, and the Bay Area Cryo-EM facility at UC Berkeley for EM imaging. We acknowledge the support of PRACE (2019215144, MG), the Marconi100 supercomputing center, and the Savio cluster at UC Berkeley for MD simulations. This work was supported by grants from the National Institute of General Medical Sciences (GM094522 (AY), GM123655-03 (LF), GM051487 (EN), GM127018 (EN) and the National Science Foundation (MCB-1617028 and MCB-1055017, AY). E.N. is a Howard Hughes Medical Institute Investigator.

## AUTHOR CONTRIBUTIONS

L.F., L.E., E.N., and A.Y. conceived the project and analyzed the data. L.F. purified the proteins, performed singlemolecule experiments, and analyzed the data. L.E. performed the cryo-EM sample preparation, data collection, and analysis. T.H. collected and analyzed data for K490. E.G. performed the kinesin pelleting assays. Q.F. improved the MT image processing pipeline and performed additional cryo-EM image analysis. A.J. assisted with cell culture and protein expression. J.F. performed and analyzed tug-of-war experiments. M. Golcuk and M. Gur performed MD simulations and analyzed the data. K.C. created scientific illustrations. L.F., L.E., E.N., M. Gur, and A.Y. wrote the manuscript, with further edits from all authors.

## METHODS

### Protein purification

Human MAP7 (UniProtKB Q14244-1), tau (UniProtKB P10636-8), and kinesin-1 (UniProtKB P33176-1) constructs have an N-terminal ZZ affinity tag, followed by a TEV protease cleavage sequence. MAP7 and tau constructs have an N-terminal ybbR sequence (DSLEFIASKLA)(Yin et al., 2005). Kinesin constructs have a C-terminal GFP and a SNAP-tag connected by a 1x GS linker sequence. The mouse BicDR1 (UniProtKB A0JNT9-1) construct has a C-terminal SNAP-tag. The BicDR1:Nanobody construct has a C-terminal GFP nanobody followed by the SNAP-tag.

Protein purification was performed as previously described (Ferro et al., 2019; Zhang et al., 2017). For MAP7 and tau constructs, a 500 mL cell pellet of Sf9 cells was thawed in 50 mL lysis buffer (25 mM HEPES pH 7.4, 1 M KCl, 10% glycerol, Roche protease inhibitor, 1 mM PMSF, 1 mM DTT, 1 mM ATP). The lysis was performed using a Wheaton glass dounce homogenizer. The lysate was spun for 45 min at 360,000 g in a Ti70 rotor. The supernatant was incubated with 1 mL IgG sepharose beads for 1 h. Beads were then washed in 50 mL lysis buffer and 50 mL storage buffer (25 mM HEPES pH 7.4, 300 mM KCl, 1 mM EGTA, 10 mM MgCl2, 10% glycerol, 1 mM DTT, 1 mM ATP). Beads were resuspended with 1 mL storage buffer and transferred to a 2 mL Eppendorf tube. For SNAP-tag labeling, the bead-bound protein was incubated with 10 nmol SNAP-ligand conjugated dye for 1 h on ice. For ybbR-tag labeling, the bead-bound protein was incubated with coenzyme-A conjugated dye and 1 μM SFP enzyme for 1 h. After labeling, beads were washed with 50 mL storage buffer. Beads were resuspended with 2 mL storage buffer and mixed with 30 μL of 2 mg/mL TEV protease for 1 h for elution. For dynein and BicDR, the same procedure was used using different lysis (50 mM HEPES pH 7.4, 100 mM NaCl, 10% glycerol, 1 mM DTT, 1 mM ATP, 2 mM PMSF, 1 Roche tablet per 50 mL) and storage (50 mM Tris pH 7.4, 150 mM KAc, 2 mM MgAc2, 1 mM EGTA, 10% glycerol, 1 mM ATP, 1 mM DTT) buffers.

K350^E236A^ and DCX expression plasmids were transformed into Rosetta2 (DE3) pLysS cells. Bacteria were grown in YT media until the culture reached an OD600 of 0.7. Cells were induced overnight at 20 °C with 0.2 M IPTG, spun down at 4,785 g for 15 min in a JLA 8.1 rotor, and combined with 50 mL lysis buffer (50 mM NaH_2_PO_4_, 250 mM NaCl, 2 mM MgCl_2_, 30 mM imidazole, 10% glycerol, 1 mM DTT, 1 mM PMSF, pH 8.0). Cells were lysed with a sonicator and spun at 117,734 g in a Ti70 rotor. The supernatant was incubated with 1 mL of Ni-NTA beads for 1 h. Beads were washed with 50 mL wash buffer A (lysis buffer without PMSF), 50 mL wash buffer B (wash buffer A plus 750 mM additional NaCl) and exchanged back into wash buffer A. Protein was eluted with elution buffer (50 mM NaH2PO4 pH 7.2, 250 mM NaCl, 1 mM MgCl_2_, 500 mM imidazole, 10% glycerol). For SFP, the same procedure was used with different lysis (20 mM Tris, 0.5 M NaCl, 5 mM imidazole, pH 8) and elution (20 mM Tris, 0.5 M NaCl, 500 mM imidazole, pH 8) buffers. Protein concentrations were determined with either Bradford assay or absorbance at 280 nm on a Nanodrop spectrophotometer. Protein preps were run on an SDS-PAGE gel to check for purity and snap-frozen in liquid nitrogen for storage.

### Microscopy

Fluorescence microscopy was performed using a Ti-E Eclipse inverted microscope body equipped with a 100x 1.49 NA Apo TIRF objective (Nikon). 488, 532, and 633 nm laser beams (Coherent) were fiber-coupled using laser-to-fiber couplers and a wave dimension multiplexer (OZ Optics). TIRF illumination was controlled using a TI-TIRF Motorized Illuminator unit (Nikon). The emission signal was filtered using 512/25, 580/40 697/75 nm bandpass filters (Chroma) mounted in a Lambda 10-B optical filter changer (Sutter). Images were collected with an electron-multiplied CCD camera (Andor, Ixon^+^). The microscope was controlled using Micromanager. The effective pixel size after magnification was 160 nm.

### Motility assays

Tubulin was purified from pig brain and labeled with biotin or fluorophores as described (Castoldi and Popov, 2003; Nicholas et al., 2014). The final percentage of labeled tubulin in the MTs was less than 5%. Coverslips coated with PEG/PEG-Biotin (Microsurfaces) were assembled into flow chambers with laser-cut Parafilm. All solutions for motility assays were in MB buffer (30 mM HEPES, 5 mM MgSO_4_, 1 mM EGTA, pH 7.0). 1 mg/mL streptavidin was added to the chamber and incubated for 2 min. Following three washes with 40 μL wash buffer (MB buffer supplemented with 0.5% Pluronic F-127, 1 mg/mL casein, 1 mM TCEP, 10 μM taxol), MTs were added to the chamber and allowed to attach for 2 min. Unattached MTs were removed by an extensive wash of the flow chamber and the chamber was exchanged into the imaging buffer (wash buffer supplemented with 150 mM KAc, 0.1% methylcellulose, 1 mM ATP, glucose oxidase, catalase, 0.4% glucose, motors, and MAPs). The sample was sealed and imaged immediately. Reported MAP concentrations represent the concentration of MAP added to the imaging buffer.

All fitting was performed in OriginPro 9 or MATLAB 2016a. Binding curves of MAP7 and tau to MTs were fit to a Hill equation 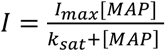, and k_sat_ was reported as half-maximal saturation, where “I” is fluorescence intensity on the MT. Motor inhibition by MAPs was fit to a single exponential decay function, where *IC_50_* is the decay rate by MAP concentration. For phase separation experiments, saturation concentration (*cSat*) was calculated from a sigmoidal logistic function, *A = A_max_*(1 + *e*^−*k*([*MAP*]−*cSat*)^)^−1^, where *A* surface area of liquid condensates, *A_max_* is the surface area of condensates at saturation and *k* is the logistic growth rate. The run frequency (*f*) of K560 was fitted to a two-component Hill equation 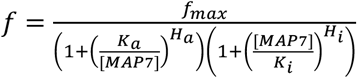, where *f_max_* is the maximum run frequency, *K_a_* is the half-maximum activating MAP7 concentration, *H_a_* is the Hill coefficient of activation, *K_i_* is the half-maximal inhibiting MAP7 concentration, and *H_i_* is the Hill coefficient of inhibition.

Partition coefficients were determined from the ratios of the fluorescence signal inside to outside the droplet. The 10-pixel square box was drawn either inside or outside the droplet. For MT binding affinity measurements, a circle was drawn to a size slightly larger than the diameter of the MT. The signal inside the box or the circle was measured with the “Measure” function in ImageJ.

### KD assemblies

The dynein complex (0.6 μM) was labeled with SNAP-LD655 on an N-terminal SNAP-tag on the dynein heavy chain and mixed with unlabeled dynactin (1 μM), and the BicDR1 construct (1 μM), which contains a C-terminal GFP-nanobody. After incubating the mixture on ice for 15 min, FL kinesin-1-GFP-SNAP labeled with LD555 (0.5 μM) was added and the mixture was incubated on ice for another 10 min. Biotinylated-MTs were attached to PEG-Biotin coverslips. Wash buffer included Pluronic and casein. The kinesin-dynein complex was diluted 500-fold in imaging buffer (wash buffer supplemented with 1 mM ATP, glucose oxidase, catalase, 0.4% glucose, and MAPs).

Movies were acquired using time-sharing between 532 and 633 nM laser exposures (200 ms each). Files were analyzed using Bio-Formats (ImageJ). In the “no MAP7” condition, immotile or diffusive spots in the dynein channel motors were attributed to dynein only motors. Stationary molecules were filtered from the kymograph in the Fourier space using a fast Fourier transform (FFT) mask in ImageJ. Two fluorescent channels were registered to identify trajectories that contain both fluorescent dyes. Velocities of these trajectories were measured by determining the distance between binding and dissociation and the time over which the motors stepped, including pauses and backtracking events.

### Sample preparation for Cryo-EM

Tubulin (Cytoskeleton) aliquots at 10 mg/ml tubulin in EM Buffer (80mM PIPES pH 6.9, 1mM EGTA, 1mM MgCl_2_, and 1 mM GTP) were polymerized for 20 min at 37° C. MTs were then centrifuged at 17,000 g for 20 min. Each MT pellet was resuspended in 20 μL EM buffer supplemented with 1 mM DTT and 266 μM peloruside (an MT-stabilizing drug). Samples were incubated at a stoichiometry of 1:2 for MTs:FL-MAP7 and 1:3:10 for MTs:MAP7-MTBD:Tau (Figure S1) for 5 min in MB buffer supplemented with 10% NP-40 and 266 μM peloruside. MT concentration was set to 6 μM. 150 mM KAc was added to the MT solution before adding FL-MAP7. The MAP/MT mixtures were added to glow-discharged C-flat holey carbon grids (CF-1.2/1.3–4C, 400 mesh, copper; Protochips) inside a Vitrobot (FEI) set at 22 °C and 100% humidity before plunge-freezing in ethane slush and then transferred to liquid nitrogen, as previously described (Eshun-Wilson et al., 2019; Kellogg et al., 2018; Zhang et al., 2015)(Figure S1).

Because MTBD MAP7 binds the MT less strongly than FL MAP7, MAP7 MTBD and FL tau were incubated with MTs in the absence of added salt.

### Cryo-EM

Micrographs were collected using an Arctica microscope (ThermoFisher) operated at an accelerating voltage of 200 kV. All cryo-EM images were recorded on a K2 Summit direct electron detector (Gatan), at a nominal magnification of 36,000×, corresponding to a calibrated pixel size of 1.14 Å. The camera was operated in superresolution mode, with a dose rate of ~1 e-per pixel s^-1^ on the detector. We used an exposure time of 4 s, corresponding to a total dose of 45 electrons/ Å^2^ on the specimen. The data were collected semiautomatically using the SerialEM software suite (Mastronarde, 2005).

### Image Processing

Stacks of dose-fractionated image frames were aligned, dose weighted, and summed using the UCSF MotionCor2 software (Zheng et al., 2017). MT segments were manually selected from 2,000 and 800 images for MAP7-FL, and MAP7-MTBD and tau datasets, respectively, using the Helical Manual Picker in RELION (Zivanov et al., 2018). We estimated the Contrast Transfer Function (CTF) using CTFFIND4 and converted the segments to 90% overlapping boxes (512 × 512 pixels) for particle extraction using the helical picking feature in RELION. The remaining non-overlapping region was set to 82 Å and corresponds to the tubulin dimer repeat (asymmetric unit). This process resulted in 63,307 and 41,021 particles for the MAP7-FL and MAP7-MTBD and tau datasets, respectively. The PF numbers of the extracted particles were sorted by performing supervised 3D classification using RELION. For both datasets, the 13-PF class showed better-resolved features. Thus, 13-PF particles, 29,993 particles for the MAP7-FL dataset, and 18,450 particles for the MAP7-MTBD-tau dataset were selected for further processing. We used RELION to determine the initial alignment parameters of the selected particles and then transferred them to FREALIGN (Grigorieff, 2016) for further refinement and for αβ-tubulin register and seam location determination of each MT segment. Pseudo-helical symmetry was imposed during 3D reconstruction using helical parameters determined using the relion_helix_toolbox function in RELION. The full microtubule reconstruction was regenerated by excising the best PF from the FREALIGN reconstruction using EMAN2 libraries, as previously described ((Alushin et al., 2014; Zhang et al., 2015). The αβ-tubulin register and seam location determination were performed according to a previously described protocol (Zhang and Nogales, 2015) that we have now automated by designing a wrapper Python script. A schematic of the MT pipeline is outlined in Figure S1. The refinement in FREALIGN gave us final maps of the MAP7-FL dataset at a resolution of 4.4 Å and of the MAP7-MTBD-tau dataset at a resolution of 4.5 Å (Figure S1). The resolution was estimated by applying a soft cylindrical mask around the MT walls.

### Co-sedimentation assays

Peloruside-stabilized MTs were incubated with combinations of MAP7 and kinesin at 25 °C for 10 min in MB buffer supplemented with 150 mM KAc, 1 mM DTT, and 2 mM peloruside. The final concentrations of MTs, kinesin, and MAP7 were 8, 16, and 24 μM, respectively. 100 μM AMP-PNP was added for stably tethering kinesin to MTs. After incubation, 20 μL reaction mixtures were centrifuged at 90,000 g for 15 min at 25 °C atop 20 μL of 50% glycerol cushion. The supernatant (20 μL) and the pellet (20 μL) were added to 5X SDS sample buffer, heated at 95 °C for 10 min, and loaded on denaturing 4-12% Bis-Tris gels (Invitrogen). The gels were stained with Coomassie blue and imaged using the Bio-Rad Gel Imager. Band intensities were determined using ImageJ and normalized to the MT pellet for each experiment.

### Atomic model building

Full-length dynein conformations were sampled from the MD simulations performed on the ADP-bound structure of human dynein-2 in the absence of MTs (Can et al., 2019). The MTBD and parts of stalk coiled-coils 1 and 2 (CC1 and CC2, respectively) of dynein-2 (amino acids 2962 – 3126) were used to investigate dynein binding to tau-free and tau-bound tubulin heterodimer. The cryo-EM structure of the R2 repeat of tau on βαβ tubulin (PDB: 6CVN)(Kellogg et al., 2018) was used as a template for simulations. For tau to span an αβ tubulin heterodimer, β-tubulin located at C-terminal of the tau structure and tau residues K298-V300 were removed, and the tau R2 structure was extended by adding residues S262-G273 to its N-terminal end. The missing region comprising residues P270-G273 was constructed using the Molefacture plugin of VMD (Humphrey et al., 1996). To define the dynein binding site on this structure, the atomic model of dynein-1 MTBD in complex with tubulin (PDB: 3J1T)(Redwine et al., 2012) was aligned with the R2-bound tubulin structure using the secondary structures of the tubulins. Subsequently, MTBD conformations sampled from our MD simulations of human dynein-2 at the low-affinity state were superimposed onto dynein-1 MTBD via helices 1, 2, 3, and 6. Finally, aligned conformations of human dynein-2 MTBD were then moved 7 Å away from the dynein binding site of the αβ tubulin.

### MD simulations

25 simulations were performed for tau-free and tau-bound conditions. Each system was solvated in a water box using the TIP3P water model with padding of at least 15 Å of water in each direction. The solution was ionized to 150 mM NaCl. Each system was composed of ~200,000 atoms. MD simulations were performed in NAMD 2.13 (Phillips et al., 2005) using the CHARMM36 all-atom additive protein force field (Best et al., 2012). A time step of 2 fs was used. For van der Waals interactions, a 12 Å cut-off distance was applied. The particle-mesh Ewald method was used to calculate long-range electrostatic interactions. The temperature was kept constant at 310 K using a damping coefficient of 1 ps^-1^ for Langevin dynamics. The pressure was maintained at 1 atm using the Langevin Nosé-Hoover method with an oscillation period of 100 fs and a damping time scale of 50 fs. For tau systems, dynein and tubulin were kept fixed for 10,000 steps of minimization, followed by 1 ns equilibration. Next, constraints on dynein and tubulin were released for an additional 10,000 steps. Subsequently, 2 ns of equilibration were performed, during which harmonic potentials having a spring constant of 1 kcal mol^-1^Å^-2^ was applied to the C_α_ atoms. Before initiating the MD simulations, all constraints were removed and the system was equilibrated for an additional 4 ns. In tau-bound simulations, harmonic constraints were applied to C_α_ atoms of tau and the tubulin residues within 5 Å of tau. The total simulation time was 12.5 μs.

Conformations sampled in MD simulations were aligned via tubulin. The x and z coordinates of the H1 center of mass were calculated. Heat maps were constructed in MatLab. Principal axes (PA) of MTs were obtained via the orient tool in VMD. PA1 corresponds to the longitudinal axis of a protofilament. PA2 (radial axis) passes through the center of masses of the Q2982 and Y3099 Cα, K2966 Cα, and E3117 Cα. PA3 (tangential axis) is perpendicular to PA1 and PA2. The stalk coiled coil vector (pointing from Q2982 and Y3099 Cα to K2966 Cα and E3117 Cα) was projected on the plane constructed by PA1 and PA2. The angle between the projected vector and PA2 was defined as the stalk angle. A similar procedure was used to calculate the angular orientation of H1. The H1 vector points from S3005 Cα to E2998 Cα. A maximum cutoff distance of 6 Å between the basic nitrogen and acidic oxygens was used to detect salt bridges (Beckstein et al., 2009). Hydrogen bonding was defined as a maximum 3.5 Å separation between the donor and acceptor atoms, together with a maximum 30° angle between the hydrogen atom, the donor heavy atom, and the acceptor heavy atom.

## Methods

### KEY RESOURCES TABLE

| REAGENT or RESOURCE | SOURCE | IDENTIFIER |
| --- | --- | --- |
| Bacterial and Virus Strains |  |  |
| Competent cells | QB3 MacroLab | XL1-Blues |
| Competent cells | QB3 MacroLab | Rosetta2 (DE3) pLysS |
| Competent cells | QB3 MacroLab | DH10Bac |
| Insect cells | Berkeley Cell Culture Facility | Sf9 |
| Chemicals, Peptides, and Recombinant Proteins |  |  |
| Glucose oxidase | Sigma | G2133 |
| Catalase | Sigma | C40 |
| Taxol | Sigma | T7191 |
| CoA Cy3 | Lumidyne | Custom synthesis |
| CoA Cy5 | Lumidyne | Custom synthesis |
| ATP | Sigma | A3377 |
| Pluronic F-127 | ThermoFisher | P6866 |
| Streptavidin | ThermoFischer | S888 |
| Methylcellulose | Sigma | M0512 |
| Casein | Sigma | C5890 |
| Taxol | Sigma | T7191 |
| TCEP | Sigma | C4706 |
| Human KIF5B (1-490 aa)::GFP:: SNAP | This study | N/A |
| Human KIF5B (1-560 aa) ::GFP:: SNAP | This study | N/A |
| Human KIF5B (1-963 aa) ::GFP:: SNAP | This study | N/A |
| Human KIF5B (1-350 aa, E236A) |  | N/A |
| Human MAP7 (1-749 aa) | This study | N/A |
| Human ybbR::MAP7 (1-749 aa) | This study | N/A |
| Human ybbR::MAP7 (1-316 aa) | This study | N/A |
| Human ybbR::MAP7 (60-170 aa) | This study | N/A |
| Human ybbR::MAP7 (307-749 aa) | This study | N/A |
| Human ybbR::Tau (1-441 aa) | This study | N/A |
| Human ybbR::Tau (242-367 aa) | This study | N/A |
| Human DCX |  | N/A |
| Porcine dynactin complex |  | N/A |
| Human dynein complex |  | N/A |
| Mouse BicDR1 (1-577 aa)::SNAP | This study | N/A |
| Mouse BicDR1 (1-577 aa)::SNAP::GFP nanobody | This study | N/A |
| Tubulin |  | N/A |
| SFP |  | N/A |
| TEV protease |  | N/A |
| Critical Commercial Assays |  |  |
| BCA assay kit | ThermoScientific | 23250 |
| Deposited Data |  |  |
| Recombinant DNA |  |  |
| Human DCX | Addgene | 83928 |
| Human KIF5B (1-490 aa)::GFP:: SNAP | This study | N/A |
| Human KIF5B (1-560 aa) ::GFP:: SNAP | This study | N/A |
| Human KIF5B (1-963 aa) ::GFP:: SNAP | This study | N/A |
| Human KIF5B (1-350 aa, E236A) |  | N/A |
| Human MAP7 (1-749 aa) | This study | N/A |
| Human ybbR::MAP7 (1-749 aa) | This study | N/A |
| Human ybbR::MAP7 (1-316 aa) | This study | N/A |
| Human ybbR::MAP7 (60-170 aa) | This study | N/A |
| Human ybbR::MAP7 (307-749 aa) | This study | N/A |
| Human ybbR::Tau (1-441 aa) | This study | N/A |
| Human ybbR::Tau (242-367 aa) | This study | N/A |
| Human dynein complex |  | N/A |
| Mouse BicDR1 (1-577 aa)::SNAP | This study | N/A |
| Mouse BicDR1 (1-577 aa)::SNAP::GFP nanobody | This study | N/A |
| Software and Algorithms |  |  |
| ImageJ | ImageJ | https://imagej.nih.gov/ij/ |
| Matlab | MathWorks | https://www.mathworks.com/ |
| Origin | OriginLab | https://www.originlab.com/ |
| BoxPlotR | BoxPlotR | http://shiny.chemgrid.org/boxplotr/ |
| Protein disorder prediction | IUPred2A | https://iupred2a.elte.hu/ |
| Protein charge distribution | EMBOSS | http://www.bioinformatics.nl/cgi-bin/emboss/charge |
| Amino acid sequence shuffle | SMS | https://www.bioinformatics.org/sms2/shuffle_protein.html |
| Other |  |  |
| Biotin PEG coverslips | Microsurfaces | Bio-02 |
| Ni-NTA beads | Thermo Scientific | 88221 |
| IgG sepharose beads | GE Healthcare | 17096902 |
| Glass dounce | Wheaton | 357544 |
| cOmplete, Mini, EDTA-free Protease Inhibitor Cocktail | Sigma | 11836170001 |

## SUPPLEMENTAL FIGURES

**Figure S1.**
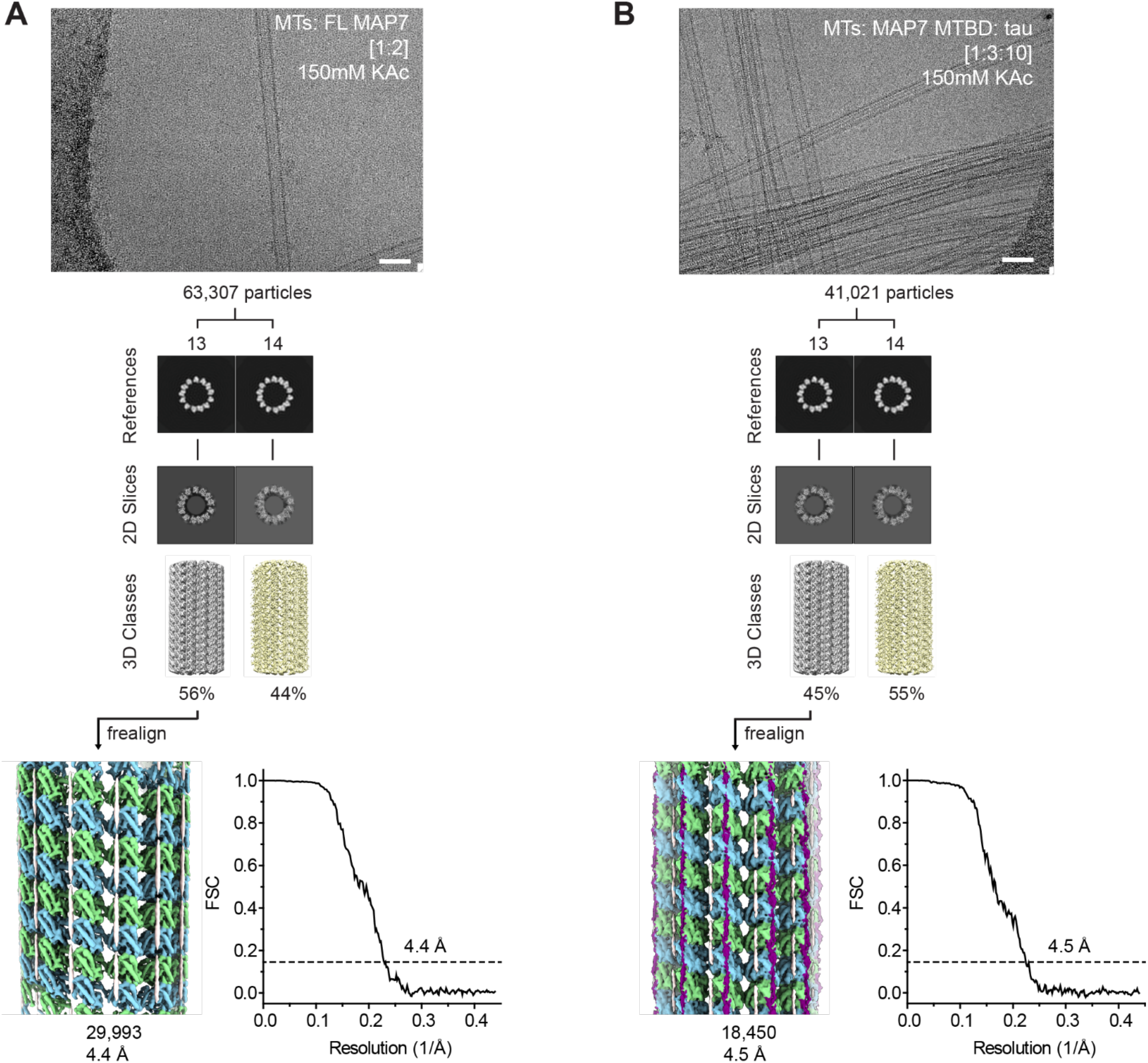
Data tree and FSC curves for the MAP7 and MAP7 in the presence of tau structures; related to Figure 1. **(A)** Representative micrographs for MTs decorated with FL MAP7. **(B)** Representative micrographs for MTs decorated with MAP7-MTBD and tau. Fourier Shell Correlation (FSC) curves for the overall maps reveal the final resolution based on the gold-standard criterion (FSC = 0.143).

**Figure S2.**
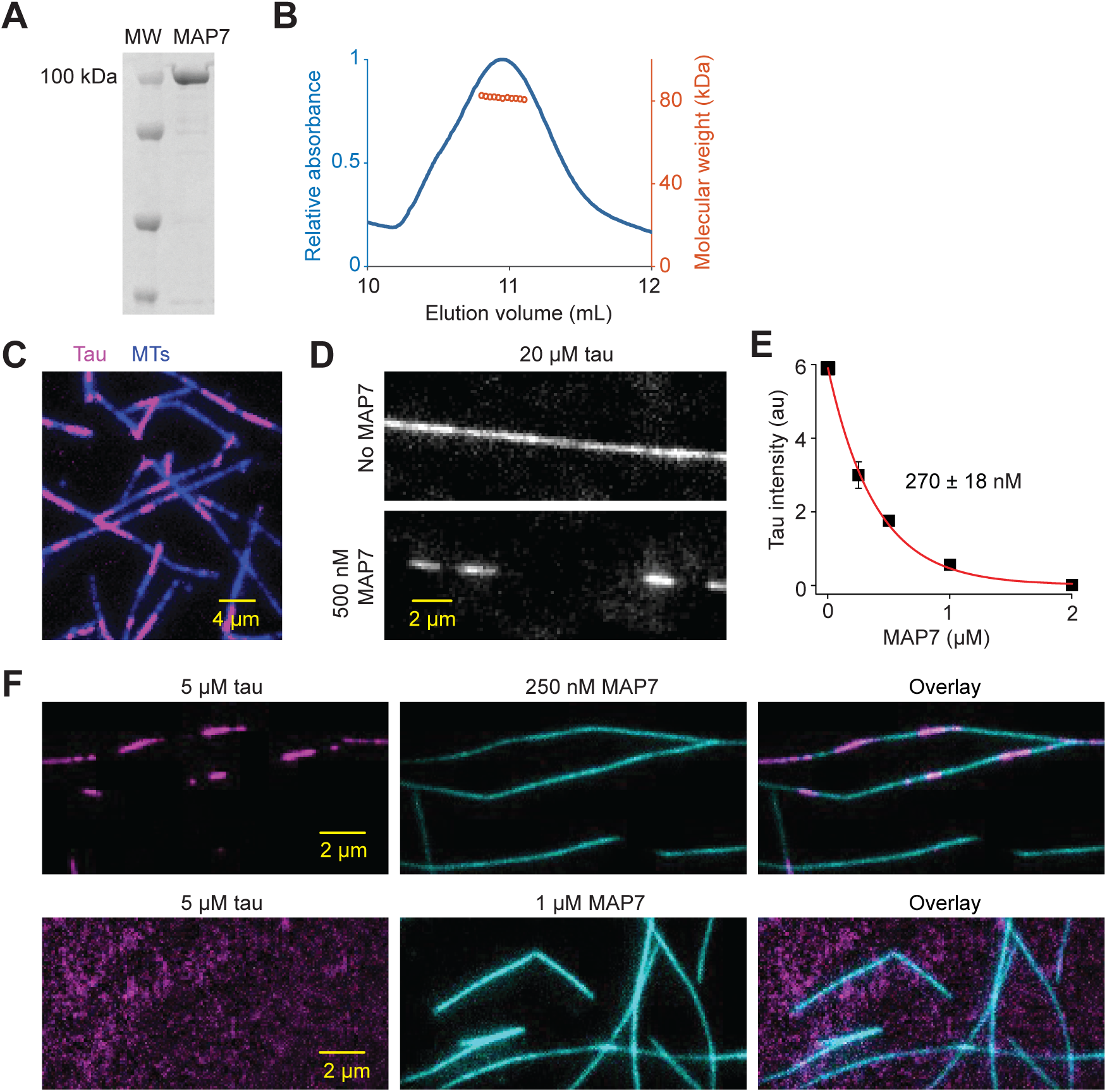
Interactions between MAP7 and tau on MTs; related to Figure 2. **(A)** Denaturing gel of purified MAP7 (MW: molecular weight). **(B)** Size exclusion chromatography coupled to the multi-angle light scattering of MAP7 (SEC-MALS) shows MAP7 elutes as a monomer. **(G)** Fluorescent image shows tau condensates on the MT. Tau concentration is 3 μM. **(H)** MT decoration of 20 μM tau in the presence and absence of 500 nM MAP7. **(I)** MAP7 removes tau from MTs. Fluorescence intensity of 20 μM tau on MTs was measured at different MAP7 concentrations (symbols, mean ± s.d.). IC_50_ (mean ± s.d.) was determined from a fit to a single exponential decay (red curve). **(J)**Two-color fluorescent images of 5 μM tau and 250 nM MAP7 show an overlap between MAP7 and tau islands on the MT. 1 μM MAP7 displaces 5 μM tau on the MT.

**Figure S3.**
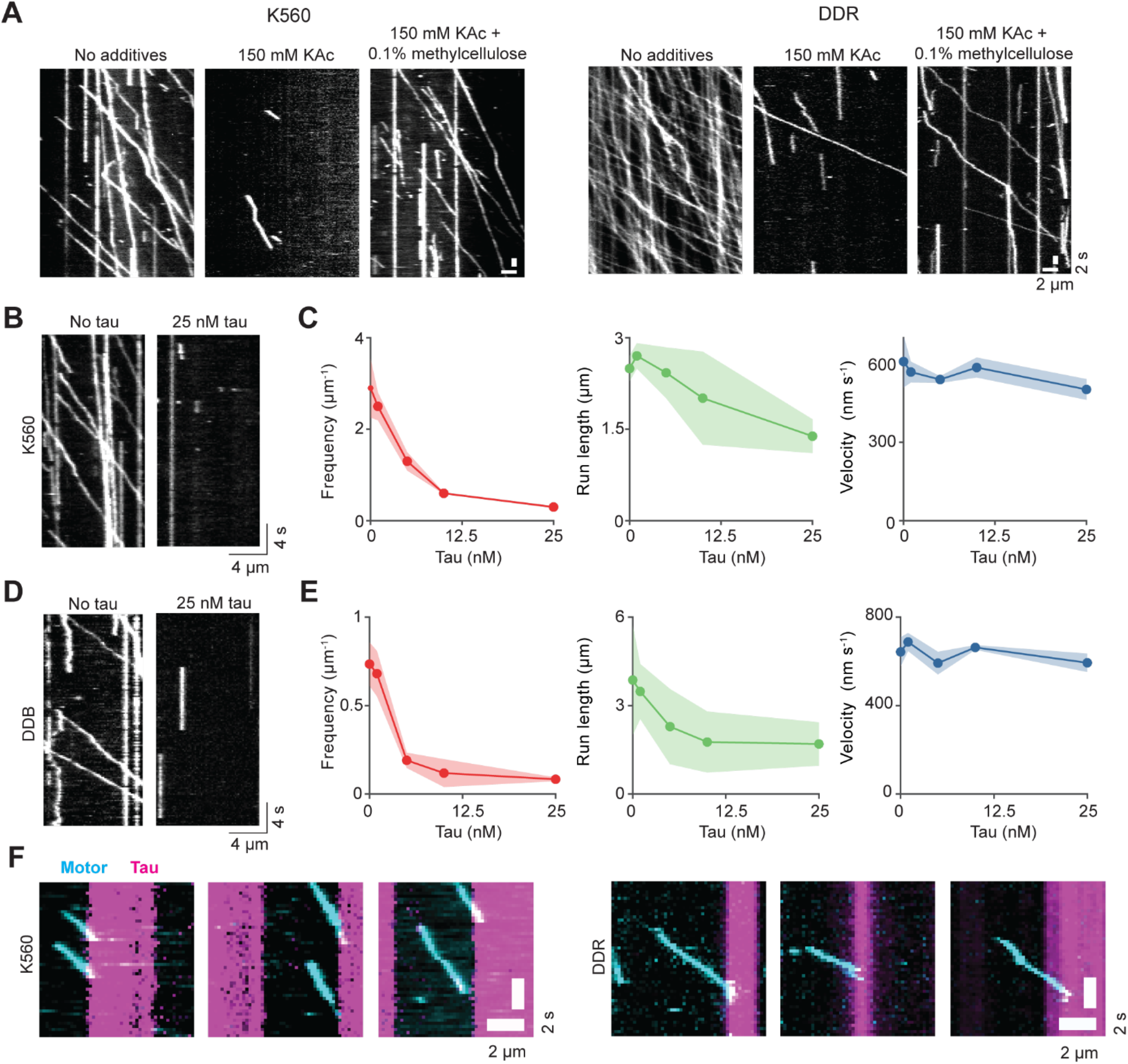
Regulation of single and a team of kinesin and dynein motors by tau; related to Figure 2. **(A)** K560 and DDR motility in the presence and absence of 150 mM KAc and 0.1% methylcellulose. **(B)** Kymographs of K560 motility with and without tau in the absence of added salt or methylcellulose. **(C)** The run frequency, run length, and velocity of K560 at different tau concentrations. N = 432, 270, 229, 193, 133 runs from left to right. **(D)** Kymographs of the dynein-dynactin-BicD2N (DDB) complex with and without tau in the absence of added salt or methylcellulose. **(E)** The run frequency, run length, and velocity of DDB at different tau concentrations. N = 210, 162, 89, 71, 50 runs from left to right. **(F)** When the motors encounter a tau condensate on an MT in 150 mM KAc, they either dissociate immediately or extend into the condensates before they quickly fall off the MT.

**Figure S4.**
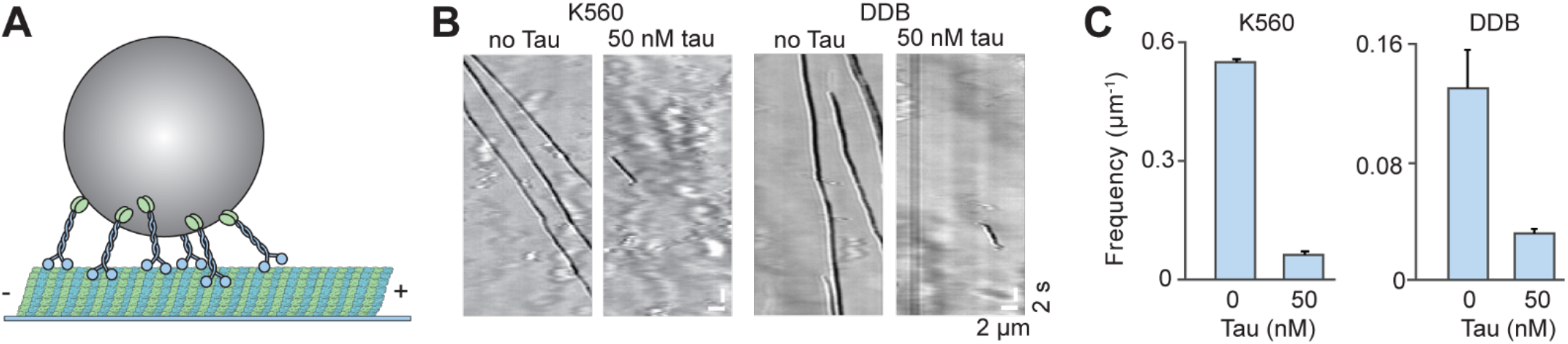
Tau inhibits beads driven by multiple kinesin motors; related to Figure 2. **(A)** Schematic shows a bead driven by multiple kinesin motors. **(B)** Beads driven by multiple K560s or the dynein-dynactin-BicD2 complexes (DDBs)(Schlager et al., 2014) are inhibited by 50 nM tau in the absence of added salt or methylcellulose. **(C)** The run frequency for multimotor driven beads decreases by tau. From left to right, N = 210, 41, 109, and 28 runs. Error bars represent s.d.

**Figure S5.**
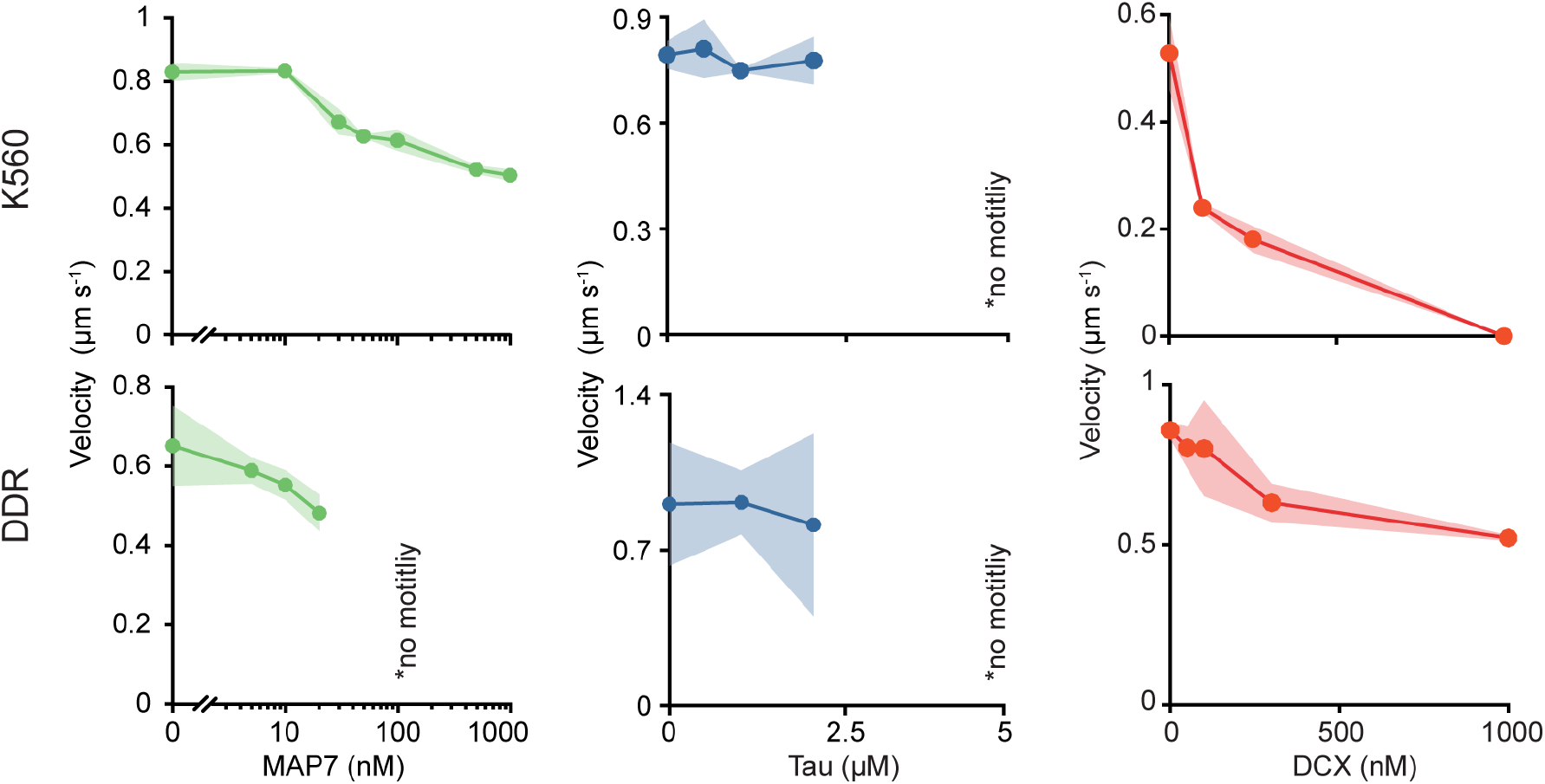
Velocity of kinesin and DDR under different MAP concentrations; related to Figure 2. Values represent mean ± s.d. For MAP7, N = 281, 463, 532, 836, 381, 433, 233 K560 motors, and 386, 235, 213, 146 DDR motors. For tau, N = 198, 160, 209, 99 K560 motors, and 185, 129, 100 DDR motors for tau. For DCX, N = 655, 187, 130, 0 K560 motors and 224, 237, 245, 104, 55 DDR motors (from left to right, two technical replicates).

**Figure S6.**
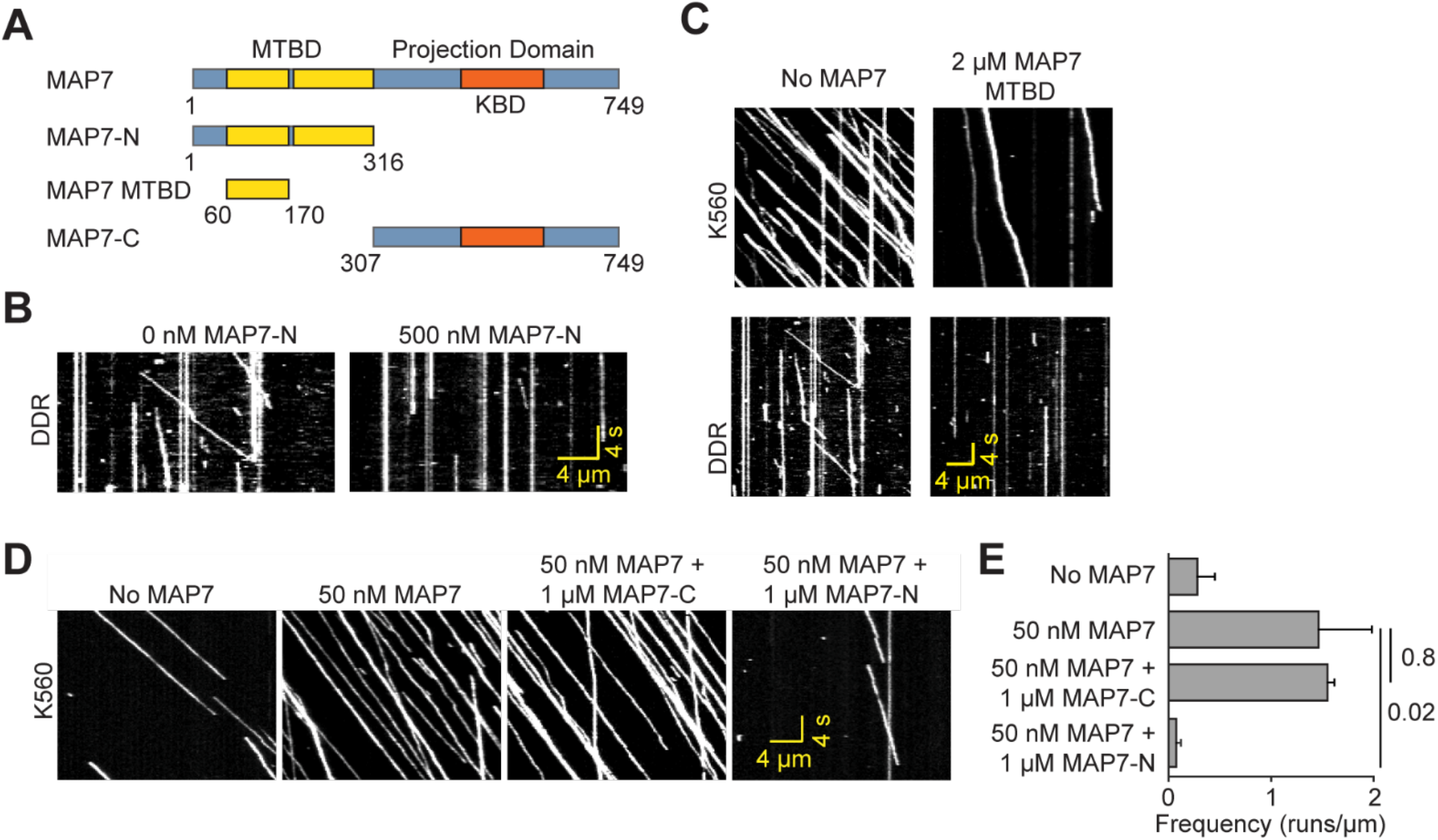
Regulation of kinesin and dynein motility by MAP7; related to Figure 3. **(A)** Schematic of MAP7 and tau truncation constructs. **(B)** Kymographs show that MAP7-N inhibits DDR motility. **(C)** Kymographs show that MAP7 MTBD inhibits K560 and DDR motility. **(D)** Kymographs of K560 motility in the presence of MAP7 construct combinations. **(E)** K560 run frequency for different MAP7 construct combinations. From top to bottom, N = 114, 355, 365, 27 runs. Error bars represent s.d. p-values are calculated from a two-tailed t-test. Assays were performed in 150 mM KAc and 0.1% methylcellulose.

**Figure S7.**
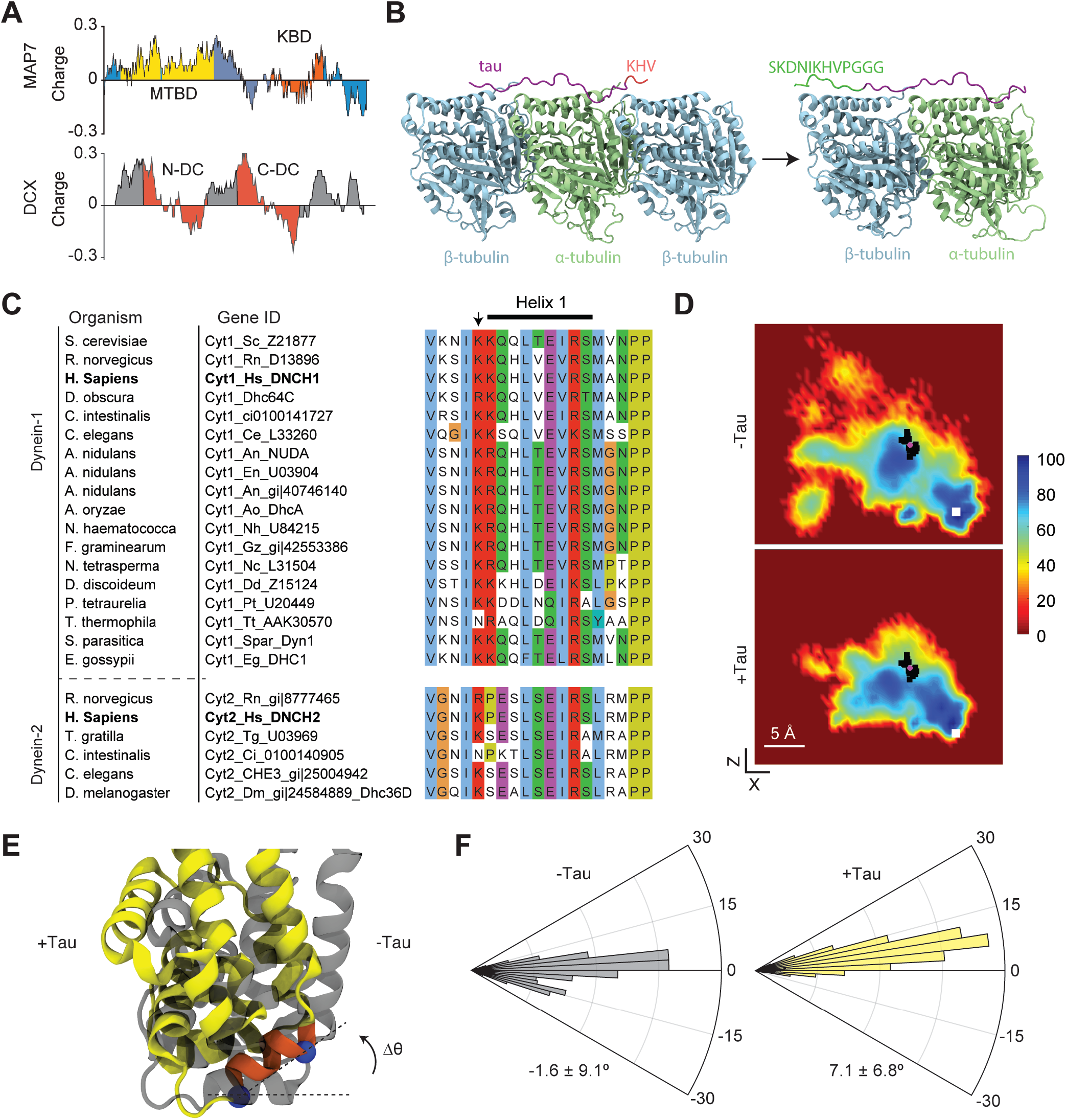
MD simulations of dynein MTBD in the presence and absence of the R2 sequence of tau; related to Figure 6. **(A)** Charge distribution on MAP7 and DCX using a sliding window size of 30 amino acids. **(B)** The structure of the R2 repeat of tau on βαβ tubulin (PDB: 6CVN) was used as a template for building the atomic model. For tau to span an αβ tubulin heterodimer, β-tubulin located at C-terminal of the tau structure and tau residues KHV were removed, and the tau R2 structure was extended by adding residues SKDNIKHVPGGG to its N-terminal end. **(C)** Alignment of dynein-1 (Cyt1) and dynein-2 (Cyt2) heavy chain amino acid sequences from different organisms. Helix 1 location is shown by the black bar above the figure. The arrow points to the conserved lysine residue (K2996 in human dynein-2 and K3298 in human dynein-1). **(D)** The heat maps show the sampling of H1 for the entire 250 ns run of 25 simulations. Black squares show the XZ position of the H1 center of mass for individual trajectories. The pink circle shows the average position of black squares. The white square shows the expected position of the H1 center of mass on tubulin at the high-affinity state of dynein. **(E)** Example snapshots of dynein MTBD in the presence and absence of tau was superimposed to show that H1 cannot retain its expected orientation on the MT in the presence of tau. Dashed lines, which pass through two well-defined positions in the H1 helix (blue spheres in the +Tau conformer), was used to quantify the angular orientation of H1. **(F)** The angular distribution of H1 (orange) in the presence and absence of tau (mean ± s.d.). The orientation of H1 in the high-affinity state of dynein (PDB: 6RZB) was set to 0°.

## SUPPLEMENTAL MOVIE LEGENDS

**Movie S1. Cryo-EM structure of MAP7 and tau on the MT**. The kinesin motor domain and the dynein MTBD were superimposed onto the MAP7 and tau structure on the MT.

**Movie S2. K560 and DDR motility on MTs in the presence of different MAP7 and tau concentrations.** K560 motors were labeled with LD555-BG on a C-terminal SNAP-tag. The DDR complex was labeled with LD555-BG on a C-terminal SNAP-tag of BicDR-1. MAP7 or Tau is added at the given concentration along with the motor solution and is not washed out from the chamber. Motility buffer includes 150 mM KAc, 0.1% methylcellulose, glucose oxidase, catalase, dextrose, and 1 mM ATP. K560 and DDR motility were recorded at 5 Hz under TIRF illumination. In a separate fluorescent channel (not shown), MT decoration on the PEG-biotin coverslip was recorded.

**Movie S3. K490 motility on MTs in the presence of different MAP7 concentrations.** K490 was labeled with LD555-BG on a C-terminal SNAP-tag. In a separate fluorescent channel (not shown), MT decoration on the PEG-biotin coverslip was recorded. Kinesin motility is shown in the presence of 0, 10, 100 nM MAP7 in the chamber. Motors and MAP7 were flown into the chamber and images were collected at 5 Hz under TIRF illumination. Motility is shown in the presence of 0, 5, 75 nM MAP7. Motility buffer includes 150 mM KAc, 0.1% methylcellulose, glucose oxidase, catalase, dextrose, and 1 mM ATP.

**Movie S4. K560 motility on MTs in the presence of different MAP7-N concentrations.** K560 motors were labeled with LD555-BG on a C-terminal SNAP-tag, and their motility was recorded at 5 Hz under TIRF illumination. In a separate fluorescent channel (not shown), MT decoration on the PEG-biotin coverslip was recorded. Kinesin motility is shown in the presence of 0, 50, 100 nM MAP7-N in the chamber. Motility buffer includes 150 mM KAc, 0.1% methylcellulose, glucose oxidase, catalase, dextrose, and 1 mM ATP.

**Movie S5. Motility of kinesin-DDR assemblies in the presence and absence of MAP7.** FL kinesin with a C-terminal GFP was labeled with LD555-BG on a C-terminal SNAP-tag. BicDR, with a C-terminal anti-GFP nanobody, was expressed in Sf9 cells. Dynein was purified from Sf9 cells and fluorescently labeled with LD655 on an N-terminal SNAP-tag. DDR complex was assembled on ice and mixed with kinesin to form kinesin-DDR assemblies. Two-color movies were collected under TIRF illumination. Each channel’s exposure time was 200 ms. The motility of assemblies is shown in the presence of 10 nM MAP7. Motility buffer includes glucose oxidase, catalase, dextrose, and 1 mM ATP.

**Movie S6. Binding of dynein to the MT in the presence and absence of the R2 repeat of tau on tubulin.** Time-lapse movies of MD trajectories show that a salt bridge between K2996 of dynein (corresponds to K3298 in human dynein-1) with E431 of β-tubulin facilitates docking of the H1 helix to its binding site. The formation of this salt bridge was interrupted by interactions between the dynein MTBD and the backbone of the R2 repeat of tau. C_α_ atoms of the tau backbone were fixed in position. The tubulin C-terminals tails were not included in MD simulations.

**Movie S7. Tau affects the orientation of dynein on the MT.** Time-lapse movies of MD trajectories show that, in the presence of tau, dynein (yellow) is tilted away from tau by ~10° in comparison to its stable docking orientation (grey) in the absence of tau.

